# A novel interoceptive subfornical organ to infralimbic cortex circuit relays airway inflammation effects on fear extinction

**DOI:** 10.1101/2025.07.16.664367

**Authors:** Emily Allgire, Jaclyn W. McAlees, Rebecca A. Ahlbrand, Elizabeth Mancz, Lauren L. Vollmer, Andrew Winter, Katherine M.J. McMurray, Laura Maile, Bryan Sanders, William G. Ryan, Allan-Hermann Pool, Igal Ifergan, Eric S. Wohleb, Steve Davidson, Robert E. McCullumsmith, Ian P. Lewkowich, Renu Sah

## Abstract

There is growing interest in the impact of internal body states on the brain and behavior. The detrimental effects of chronic lung inflammation on mental health are well recognized, however, underlying mechanisms are not known. Here, using a murine model of allergic asthma we report compromised fear extinction in mice with severe but not mild airway inflammation (AI); an effect abolished by anti-interleukin-17A (IL-17A) antibodies. Investigation of innate immune cells, microglia as-well-as transcriptomic signatures in the subfornical organ (SFO), a brain interoceptive node lacking a traditional blood-brain-barrier, revealed significant alterations in severe AI mice. IL-17 Receptor A (IL-17RA) was expressed in SFO microglia and upregulated in severe AI mice. Notably, ablation of microglial IL-17RA improved fear extinction in severe AI mice. Furthermore, we identified direct SFO projections to the infralimbic (IL) cortex, a key area regulating extinction. Importantly, chemogenetic inhibition of the SFO-IL circuit led to improved fear extinction in severe AI mice. Collectively, we report a unique body-to-brain interoceptive mechanism engaging the SFO and an SFO-to-IL circuit, through which airway inflammatory mediators compromise fear extinction. Beyond asthma, our findings are relevant to other pulmonary pathologies (e.g. bacterial pneumonia, ARDS, COVID-19) highlighting a risk for cortical dysfunction and fear pathologies such as PTSD.

## Introduction

In recent years there has been an appreciation of the adverse effects of inflammation within peripheral organ systems such as the lungs on mental health^1^; although underlying mechanisms are not understood. Strong evidence supports a high prevalence of psychiatric illnesses in individuals with a history of severe airway inflammation^2^, as observed in acute respiratory distress syndrome, ARDS^3^, COVID-19^4,5^, and chronic asthma ^6^. More than 300 million people globally, most notably children, suffer from allergic asthma^7,8^, a chronic inflammatory lung condition. Asthma has been associated with depression and anxiety ^9,10^, as well as posttraumatic stress disorder (PTSD) ^11^, a disorder of persistent fear memories. In fact, a strong genetic correlation (r_g_ = 0.49) exists between asthma and PTSD ^12^, and in a cohort of veterans with PTSD screened for 28 chronic physical illnesses the highest odds ratio was reported for asthma (OR= 8.26)^13^. This relationship seems to be stronger in severe asthmatics as opposed to mild/moderate asthmatics^14,15^. Despite this knowledge, the association between asthma-relevant chronic airway inflammation and emotional fear responses remains unclear.

Persistence of fear in PTSD results from impaired fear extinction^16^. The medial prefrontal cortex (mPFC) has been identified as a key area regulating fear extinction^17^. Reduced mPFC activity, particularly in the ventromedial infralimbic (IL) subdivision, is associated with compromised fear extinction in individuals with PTSD^16^. Emerging evidence shows that cortical function is modulated by asthma-related airway inflammation ^18–19^ However, the exact mechanisms underlying these associations are currently unknown. Identification of afferent mechanisms that can sense and relay the effects of asthma-associated airway inflammation to the IL for fear regulation is therefore highly relevant.

Circumventricular organs (CVOs), regarded as “windows on the brain”, are strategic interoceptive nodes localized adjacent to the ventricles ^20^. Fenestrated capillaries within CVOs result in a compromised blood brain barrier (BBB) that enables sensing of the internal state of the body for homeostatic maintenance^21^. Notably, sensory CVOs, such as the subfornical organ (SFO), organum vasculosum of the lamina terminalis (OVLT) and area postrema (AP) possess strategic neural connections to forebrain effector sites enabling integration of systemic cues with brain function and behavior^22^. Among sensory CVOs, the SFO has gained attention as a receptive locus for adaptive immune cells ^23^ and is known to regulate the central actions of circulating cytokines such as IL-6, IL-1β, and TNFα ^24–26^. Thus, in addition to its well-recognized role in the homeostatic control of thirst and salt appetite ^27,28^, autonomic balance^29^ and energy metabolism^30^, the SFO serves as an important interoceptive portal for sensing and relaying the effects of systemic inflammatory mediators. However, whether the SFO can sense and translate the effects of asthma-associated inflammatory mediators on cortical regulation of fear is not known.

To fill these knowledge gaps, we utilized a murine house dust mite (HDM) model of allergic asthma able to simulate heterogeneity in severity of airway dysfunction (i.e. severe vs mild) to interrogate mechanistic links between airway inflammation and fear extinction. Mild asthma is associated with increased T helper 2 (Th2) cell driven eosinophil recruitment, airway remodeling, and bronchoconstriction ^31^. However, in more severe forms of asthma this phenotype is accompanied by the additional expansion of T Helper 17 (Th17) cell driven neutrophilic responses and production of cytokines such as interleukin-17A (IL-17A) that exacerbate the symptoms of asthma^32^. Currently, cell-circuit mechanisms linking airway inflammation-associated effects and IL-mediated fear have not been established.

Here, using complementary behavioral, transcriptomics, transgenics, FACS, and intersectional chemogenetic approaches we demonstrate a key role of SFO microglia and SFO-to-infralimbic cortical projections in driving allergen induced IL-17A-mediated deficits in fear extinction. These novel observations highlight a unique role of a circumventricular organ, the SFO, as an afferent sensory node that can sense and directly relay organismal information to the cortex for emotional regulation.

## Methods

### 1. Animals

The University of Cincinnati (UC) Animal Care and Use Program (ACUP) encompasses Laboratory Animal Medical Services (LAMS, animal facilities). Some studies were carried out at the Cincinnati Children’s Hospital Research Foundation. Both are AAALAC accredited animal facilities, and all animal protocols were approved by institutional IACUCs. Adult male BALB/c mice were ordered from Jackson Laboratories (Bar Harbor, Maine, USA) at 7 weeks. The CreloxP recombination system was utilized to achieve cell-type specific, constitutive knockout of the interleukin 17 receptor A (*Il17ra*) gene. The Cx3cr1^CreER^ (JAX: 020940) transgenic mouse line in which the expression of Cre recombinase is under the control of the *Cx3cr1* promoter was crossed with the floxed *Il17ra* mouse line B6.Cg-*Il17ra^tm2.1Koll^*/J (JAX: 031000). For assessment of Fos expression, B6.129(Cg)-Fos^tm1.1(cre/ERT2^/J (JAX: 021882) were crossed with B6.Cg-Gt(ROSA)26Sor^tm9(CAG–tdTomato)Hze^/J (JAX: 007909, Ai9 flox) for fluorescent detection of “TRAP”ed cells upon Cre recombination. Mice were pair-housed in a climate-controlled vivarium with (temperature averages 23 ± 4 °C, humidity averages 30 ± 6%), ad libitum access to food and water, and maintained on a 12/12 light/dark cycle (6:00 am-6:00 pm). All housing conditions and procedures were in compliance with the National Institutes of Health Guidelines for the Care and Use of Animals. All experiments were performed between 10-14 weeks of age. Separate cohorts were used for behavioral studies and behavior-naïve animals were used for IHC, flow cytometry, FACS and RNAseq endpoints.

### 2. Treatment Protocols

#### 2.1 Aeroallergen House Dust Mite (HDM) exposure

To stimulate mild versus severe AI phenotypes we used aeroallergen house dust mite (HDM) ^33^. In brief, following acclimation to the colony, BALBc mice were given 40 µl of 100 µg of HDM (Lot 394844 with 195 EU Endotoxin/mg of protein; Greer Labs, Lenoir, NC) or phosphate buffered saline (PBS) intratracheally while anesthetized with isoflurane. HDM treatments were weekly for a total of 3 weeks (days 2, 9, 16) (see Fig 1A, B). To generate severe versus mild airway hyperresponsiveness and related immune responses, one day prior to each HDM treatment (days 1, 8, 15) mice were intratracheally administered either 35 µg αC5aR1 (BioLegend clone 20/70) or 35 µg isotype control antibody Rat IgG2b,κ (BioLegend) as described previously^33^ (HDM-αC5aR1: severe and HDM-IgG: mild). Control groups received PBS/IgG (control) or PBS/ αC5aR1 (antibody control). To assess the necessity of IL-17A in the behavioral effects of HDM exposure, animals underwent HDM/αC5aR1 treatment as described, with intraperitoneal (IP) administration of αIL-17A antibody (BioXCell) or isotype control IgG1 (BioXCell) one day prior to and after HDM treatment. (For treatment layouts and timeline, refer to schematics in figures.)

**Figure 1:**
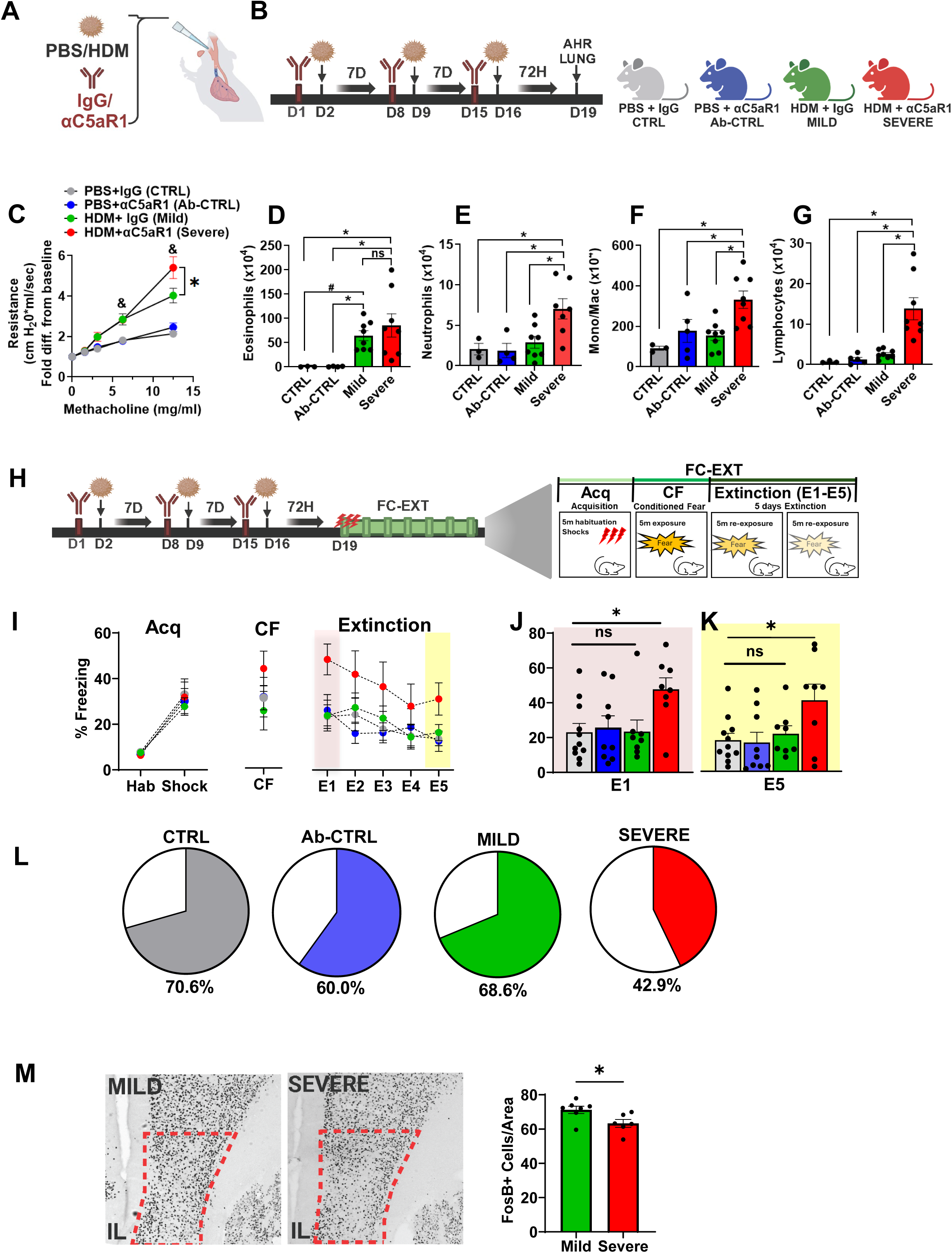
Impaired extinction of fear in mice with allergen House Dust Mite (HDM)-induced severe but not mild airway inflammation (AI): reduced neuronal activation in the infralimbic cortex of severe AI mice. (A) Intratracheal administration of PBS/HDM +/− IgG or anti-complement C5a receptor R1 antibody (αC5aR1) (B) Schematics and timeline of the experimental layout and experimental groups: PBS+IgG (CTRL), PBS+αC5aR1 (antibody, Ab-CTRL), HDM+IgG (mild) and HDM+ αC5aR1 (severe) (C) Airway hyperresponsiveness in HDM-IgG exposed mice (mild) is exacerbated in HDM+ αC5aR1 treated mice (severe) compared to PBS and αC5aR1 control groups [2-way ANOVA: treatment F _3, 78_ =16.14, p<0.0001, methacholine dose F _4,78_ =52.85, p<0.0001, interaction F_12, 78_= 5.883, p<0.0001; Significantly higher responsiveness in severe AI versus mild AI mice at 12.5mg methacholine (p=0.0001) [N=3 (CTRL),5 (Ab-CTRL), 8 (mild), 8 (severe), (D) Bronchoalveolar lavage fluid (BALF) eosinophils are increased in mild and severe AI mice compared to controls (1-way ANOVA: F _3,19_ = 4.781; p= 0.012). Significantly higher eosinophils in HDM treated mild and severe mice compared to controls (post hoc analysis: severe-CTRL p=0.011, severe-Ab-CTRL p=0.005, mild-CTRL p=0.048, mild-Ab-CTRL p= 0.044). No significant differences between mild vs severe AI (p>0.05). (E) BALF neutrophil (F) monocyte/macrophage and (G) lymphocyte counts are significantly higher in severe AI mice compared to mild, Ab-CTRL and CTRL groups. [neutrophils: 1-way ANOVA: F _3,18_ = 5.767; p= 0.006, post hoc analysis: severe-mild p= 0.006, severe-CTRL p=0.009, severe-Ab-CTRL p= 0.006; monos/macros: 1-way ANOVA: F _3,20_ = 6.26; p= 0.003, severe-mild p=0.003, severe-CTRL p=0.0035, severe-Ab-CTRL p= 0.014; lymphocytes: 1-way ANOVA: F _3,20_ = 11.85; p= 0.0001, severe-mild p=0.0002, severe-CTRL p= 0.0005, severe-Ab-CTRL p= 0.0002] No significant differences were observed between mild, CTRL and Ab-CTRL groups (p>0.05). (H) Schematic of behavioral testing for fear-conditioning-extinction (FC-EXT) in mice with mild or severe AI compared with CTRL and Ab-CTRL (I) No significant group differences in percent freezing were observed during day 1 fear acquisition {RM-ANOVA time (F_1,32_ = 72.15; p < 0.0001) treatment (F_3,32_ = 0.1294; p = 0.9418), or interaction (F_3,32_ = 0.1885; p =0.903). N= 11 (CTRL), 9 (Ab-CTRL), 8 (Mild), 8 (Severe). Conditioned fear (CF) 24 h later showed no difference in freezing between groups (F_3,32_ = 0.1595; p = 0.922). Fear extinction over 5 days revealed effects of time (F _3, 96_ = 9.009; p= 0.0001), treatment (F _3,32_ = 2.408; p = 0.0853) but no interaction (F _12,128_ = 0.9857; p = 0.4661). (J) Freezing during early extinction learning (E1) showed significant overall effect of treatment, 1-way ANOVA (F _3,32_ = 3.393; p = 0.0297). Post hoc analysis showed significantly higher freezing in severe AI mice (severe-CTRL p=0.0208, severe-Ab-CTRL p=0.022, severe-mild AI p= 0.0218). No significant differences between mild AI, CTRL, Ab-CTRL groups (p>0.05). (K) Late extinction retrieval (E5) freezing showed a significant treatment effect (F_3,32_ = 3.352; p = 0.031). Post hoc analysis revealed significantly higher freezing in severe AI mice (severe AI-CTRL (p=0.019), severe AI-Ab-CTRL (p=0.02) severe AI-mild AI (p= 0.054). No significant differences in freezing were observed between mild AI, CTRL, Ab-CTRL groups p>0.05. (L) Pie charts showing the percent distribution of mice within each group that achieved extinction (defined as <10% freezing on E5). (M) Post behavior ΔFosB cell counts within the infralimbic cortex (IL) showed a significant reduction in severe AI versus mild AI mice (t test; t_11_ = 2.48; p = 0.030, N= 7 (mild), 6 (severe)). All data are mean ± sem. * p<0.05 between groups as indicated; & p<0.05 between mild or severe AI with CTRL and Ab-CTRL groups (panel C).

#### 2.2 Tamoxifen (TAM) and 4-Hydroxytamoxifen (4-OHT) administration

TAM injections were administered based on mouse body weight (BW). Young adult *Cx3cr^cre^-Il17ra* flox mice were given two injections of TAM (100 mg/kg, IP) 48 h apart at 6-7 weeks of age. TAM powder (Sigma-Aldrich) was dissolved in sunflower seed oil (Sigma-Aldrich) to make a 20 mg/ml solution. Intratracheal administration of HDM began two weeks following the last TAM injection to allow for repopulation of IL-17RA expressing peripheral macrophages. To assess Fos-induced activation of Cre/ERT2 transgene, 50 mg/kg 4-OHT was injected intraperitoneally in B6.129(cq)-Fos^tm1.1(cre/ERT2)^ / J-Ai9 mice following SFO targeted administration of aCSF or IL-17A.

### 3. Airway hyperresponsiveness measurements

The flexiVent system (SCIREQ Scientific Respirator Equipment, Inc, Montreal, Quebec, Canada) was utilized to evaluate airway hyperresponsiveness 72 h after final HDM or PBS exposure. Mice were anesthetized with a mixture of ketamine (90–120 mg/kg), xylazine (10–20 mg/kg) and paralyzed with pancuronium bromide (0.8–1.2 mg/kg). Tracheas were cannulated with an 18-gauge blunt cannula. Mice were ventilated at 150 breaths/min, 3.0 cm water positive and expiratory pressure, and allowed to stabilize on the machine for 2 min. Mice were then exposed to methacholine (0, 3.125, 6.25, and 12.5 mg/ml) aerosolized in PBS for 15s and ventilated for an additional 10s. Ventilation cycle measurements were taken until resistance peaked. Airways were then re-recruited by deep inflation and the next methacholine dose was administered.

### 4. Behavioral Studies

#### 4.1 Contextual Fear Conditioning and Extinction

Behavioral testing for contextual fear conditioning and extinction was conducted at 72 h after final allergen administration, based on studies reporting peak airway inflammation post-HDM^34^ without effects on locomotor activity^35^. Sound attenuated isolation cabinets housing operant chambers with stainless-steel grid bars that delivered scrambled electric shocks were used (CleverSys Inc.). For all manipulations, the chambers were cleaned with 10% ethanol and allowed to dry between subjects. Fear acquisition (Acq), conditioned fear (CF) and extinction (Ext) were investigated. In brief, for acquisition mice were placed in operant chambers and allowed to habituate for 5 min and then received 3 consecutive foot shocks spaced 1 min apart (0.5 mA intensity, 1s duration). The next day, animals were placed in the chamber again for 5 min with no shocks to assess conditioned fear. For the next 5 consecutive days, mice were exposed to the chamber and recorded for 5 min without shocks to measure extinction of fear. Freezing behavior defined as complete lack of movement except respiration was measured using FreezeScan software (CleverSys Inc.) for all days. Freezing on extinction day 1 and extinction day 5 was analyzed as a measure of extinction learning (E1) and retrieval (E5), respectively.

#### 4.2 Other Behaviors

To screen for exploratory and defensive behaviors relevant to anxiety, panic and depression, a separate cohort of PBS/HDM/IgG2/ αC5aR1 exposed mice were tested on the elevated zero maze (EZM), carbon dioxide exposure (CO_2_) and forced swim test (FST).

##### 4.2.1 Elevated Zero Maze (EZM)

Performance on the EZM testing was performed using a maze (Stoelting Co., Wood Dale, IL) with four quadrants on a raised circle. Opposing quadrants consisted either of 1 cm high clear acrylic curbs or black acrylic walls 20 cm in height, characterized as the open and closed quadrants, respectively. Behavior was conducted under dim halogen lighting (24 lx on the open quadrant). Mice were initially placed in the closed quadrant and exploratory activity was video recorded for 5 minutes. Between mice, the apparatus was cleaned with 10% ethanol. TopScan software (CleverSys. Inc., Reston, VA) scored time in open and closed quadrants, latency to enter open quadrant, and total distance traveled.

##### 4.2.2 Forced Swim Test (FST)

Forced Swim Test was performed as previously described ^36^. In brief, glass cylinders (14 cm diameter, 19 cm high) were filled with water to 14cm at 24°C ± 0.5. Mice were placed in the cylinder and recorded for 6 min. Immobility (no active effort to swim or struggle) and latency to immobility were quantified by a blind observer. Average immobility was quantified as total time spent immobile from minutes 2-6.

##### 4.2.3 Carbon Dioxide (CO_2_) Inhalation

Inhalation of CO_2_ is an intense interoceptive stimulus that produces panic attacks in humans^37^. CO_2_-evoked freezing in rodents has been used as a panic-relevant behavioral assessment ^38–40^. Mice were placed in the bottom compartment of a dual vertical Plexiglas chamber (25.5 cm x 29 cm x 28 cm per chamber) and allowed to habituate for 7 min. After 24h, animals were placed back in the chamber while CO_2_ (5%; custom industrial mix in breathing air, Linde Gas & Equipment Inc., Cincinnati, OH) filled the upper chamber at an infusion rate of 10L/min. CO_2_ concentration in the lower chamber was measured by CARBOCAP® GM70 carbon dioxide meter (GMP221 probe with accuracy specification ± 0.5%, Vaisala, Helsinki, Finland) and the ambient CO_2_ concentration was confirmed to be 5.0 ± 0.5%. Mice were left in the lower chamber with CO_2_ for 10 minutes. On the next day, the animals were placed in the lower chamber in the absence of CO_2_ for five minutes to measure context conditioned freezing. On all days, freezing (the lack of all movement except respiration), was measured as a fear-relevant output utilizing FreezeScan software (CleverSys Inc., Reston, VA).

### 5. Tissue preparation and flow cytometry

#### 5.1 Tissue Collection

Lung and brain tissue was collected for flow cytometric analyses at 72 h after the final allergen administration. This time point was selected as it represents the peak post HDM inflammatory response in the lung^34^. Whole brain analysis was conducted to identify broad changes in immune composition that may have occurred in mild and severe AI mice.

Lungs were removed, minced and placed in 6 ml of RPMI 1640 containing Liberase CI (0.5 mg/ml) (Roche Diagnostics) and DNase I (0.5 mg/ml) (Sigma) at 37 °C for 45 min. The remaining tissue was forced through a 70-µm cell strainer, and red blood cells were lysed with ACK lysis buffer (Thermo Fisher Scientific). Cells were washed with RPMI containing 10% FBS, viable cells were counted via trypan blue exclusion.

Brain tissues were harvested to isolate mononuclear cells as previously described (PMID: 38906887), with modifications. Briefly, mice were anesthetized and perfused transcardially with ice-cold 1X HBSS lacking Ca²⁺ and Mg²⁺. Brains were minced with a scalpel and passed through a 100 µm filter, then subjected to enzymatic digestion using a cocktail containing 50 µg/mL DNase I and 500 µg/mL Collagenase II (Sigma). Following dissociation, cells were resuspended in a 20% Bovine Serum Albumin solution and centrifuged at 1,000 × g for 10 min to remove excess myelin and debris.

#### 5.2 Flow cytometry

Single cell suspensions of lung and brain tissue were first incubated with Fc Block (mAb clone 2.4G2) to prevent non-specific Ab binding for 15 min. Following this, cells were incubated with fluorochrome-labeled antibodies to surface markers and incubated with a fixable Live-Dead Dye to exclude dead cells (Thermo Fisher Scientific). Cells were then fixed and permeabilized for 1 h using Foxp3 staining kit (Thermo Fisher Scientific). Intracellular Fc receptors were blocked using Fc Block suspended in Permeabilization Buffer (Thermo Fisher Scientific), followed by staining with cytokine- or transcription factor-specific mAbs suspended in Permeabilization Buffer. Data were acquired with an LSR-Fortessa flow cytometer equipped with lasers tuned to 355 nm, 405 nm, 488 nm, 561 nm, and 640 nm, and digital DiVa Software. Spectral overlap was compensated, and data was analyzed using FlowJo software (Treestar Inc., Ashland, OR). All staining reagents used were purchased from Thermo Fisher Scientific, unless otherwise indicated. Clones used to stain lung cells were as follows: BV605-conjugated anti-mouse CD90.2 (clone 53–2.1, BioLegend), APC-Cy7-conjugated anti-mouse TCRβ (clone H57-597), AlexaFluor700-conjugated anti-mouse CD3ε (clone eBio500A2), Brilliant Violet 711-conjugated anti-mouse CD4 (clone RM4-5, BioLegend); PE-eFluor-610-conjugated anti-mouse CD44 (clone IM7), PE-Cy7-conjugated anti-mouse TCRγ/δ (clone GL3, BioLegend); Brillian Violet 421-conjugated anti-mouse CD8a (clone 2G12, BioLegend); Fixable Live/Dead e506; PE-conjugated anti-mouse IL-17A (clone eBio17B7), eFluor 660-conjugated anti-mouse IL-13 (clone eBio13A). Additional clones used to stain brain cells were as follows: Brilliant Violet 510-conjugated anti-mouse CD45 (clone 30-F11, BioLegend); AlexaFluor700-conjugated anti-mouse CD3ε (clone 17A2, BioLegend); Brilliant Violet 421-conjugated anti-mouse CD11b (clone M1/70, BioLegend); APC-Cy7-conjugated anti-mouse CD44 (clone IM7, BioLegend); PerCP-Cy5.5-conjugated anti-mouse CD4 (clone GK1.5, BioLegend); Brilliant Violet 786-conjugated anti-mouse CD8 (clone 53-6.7, BD Biosciences).

#### 5.3 Bronchoalveolar lavage

A cannula was inserted through the trachea and lungs were washed three times with 1 ml Hank’s Balanced Salt Solution (HBSS, Gibco 14025076) to flush airways. Collected fluid and cells were placed into a 1.5 ml centrifuge tube and centrifuged at 300 × *g* for 6 min. Fluid volume was recorded and removed (stored at −80°C) and the cell pellet was resuspended in 150 ul ACK lysis buffer (Thermo Fisher Scientific) to lyse red blood cells and incubated at room temperature for 4 min. 500 ul of PBS with 10% FBS was added to the cells to neutralize ACK and then the tubes were spun at 300 × *g* for 6 min. Supernatant was aspirated, cells were resuspended in 500 ul PBS with 10% FBS and kept on ice during total cell counting with a Hemacytometer. Approximately 50,000 cells were put onto a slide using a cytospin and then differentially stained using the DiffQuik stain kit (Camco, product#702). Cells were then differentially enumerated as either monocytes/macrophages, eosinophils, neutrophils, lymphocytes, or epithelial cells.

### 6. c-Fos reporter activation

4-hydroxytamoxifen (4-OHT, Sigma Aldrich, H6278) was dissolved in sunflower seed oil at a concentration of 10 mg/ml.-To determine the number of “TRAPed” cells, B6.129(cq)-Fos^tm1.1(cre/ERT2)^ / J-Ai9 floxed mice were injected with 25 ng of IL-17A (eBioscience) or artificial cerebrospinal fluid (aCSF) targeted to the SFO and one hour later were given an intraperitoneal injection of 50 mg/kg 4-OHT. Seven days after injection, mice were perfused transcardially with 4% paraformaldehyde and brains removed. Tissue was sectioned at 30 µm on a microtome, placed on slides and coverslipped. Images were captured and tdTomato+ cells were counted in the SFO.

### 7 Microglial gene expression

#### 7.1 Percoll enrichment of single cells

At 3 h after the last HDM treatment mice were perfused with PBS. SFO and PFC enriched punches were dissected and placed in ice-cold 1× PBS. Samples were gently homogenized via pipette mixing to minimize post-extraction disruption of microglia. Homogenized samples were centrifuged at 1200×*g* for 5 min. Supernatants were removed and cell pellets were re-suspended in 30% Percoll (GE Healthcare, Uppsala, Sweden, #17089102) and PBS. The suspension was centrifuged at 2000 x g for 20 min. The supernatant and fatty layers were removed and cells were washed with FACS buffer.

#### 7.2 Fluorescence-activated cell sorting (FACS)

Following Percoll enrichment, samples were stained for surface antigens, as previously described^41^ with modifications. In brief, Fc receptors were blocked with anti-CD16/CD32 antibody (BioLegend) for 5 min. Cells were washed and then incubated with conjugated antibodies (PE-P2RY12, BioLegend; PerCP-Cyanine5.5-CD11b, BioLegend) for 1h at 4^0^C. Cells were washed and then re-suspended in FACS buffer for analysis. Non-specific binding was assessed using isotype-matched antibodies. Antigen expression was determined using a BioRad S3e four-color cytometer/cell sorter. Cells were checked for doublets and CD11b+/CX3CR1+ population was sorted. Other myeloid cells (i.e., macrophages/monocytes: CD11b^+^/P2RY12^−^) were excluded from the sorted sample. Data were analyzed using FlowJo software (Ashland, OR, U.S.A.).

#### 7.3 Quantitative PCR

mRNA from SFO and prefrontal samples was extracted using Norgen Single Cell RNA Purification Kit (#51800) according to manufacturer protocol. In brief, cells were lysed, and DNA was removed using RNAse-Free DNase I Kit (#25710). Columns were washed with wash solution and final RNA was extracted via elution. mRNA samples were reverse-transcribed to cDNA using High-Capacity cDNA RT Kit (ThermoFisher, #4368814) according to manufacturer protocol. cDNA was diluted 1:8 and qPCR was conducted using SYBR Green Supermix (BioRad #1725121). Samples were run in duplicate for IL-17Ra and Gapdh gene expression on the QuantStudio 5 384-well PCR machine (Applied Biosystems).

### 8. Immunohistochemistry

Animals were perfused via 4% paraformaldehyde (PFA) at desired timepoints. For pre-behavior microglial staining tissue was collected a 72 h post HDM. For post-behavior ΔFosB assessment mice were sacrificed 24 h after the last behavioral measurement. Brains were extracted and submerged in 4% PFA for 24 h followed by 30% sucrose. Brains were sliced at 30µm via sliding microtome and stored in cryoprotectant (0.1M phosphate buffer, 30% sucrose, 1% polyvinylpyrrolidone, and 30% ethylene glycol) at −20°C. Slices were rinsed in 1x PBS (pH 7.4; 40 mM potassium phosphate dibasic, 10 mM potassium phosphate monobasic, and 0.9% sodium chloride) 5×5 min each. Sections were washed in 0.3% H_2_O_2_ for 10 min. Following another 5 x 5 min round of rinses, tissue was incubated in blocking solution for 1 h at RT (0.5% bovine serum albumin (BSA) and 0.4% Triton X-100). After blocking, tissue was immersed overnight in primary antibody, FosB, (1:20,000 Abcam #ab184938) or ionized calcium-binding adapter molecule (Iba-1; 1:1000 Synaptic Systems #234003). The following day, sections were washed again (5×5 min) in PBS. For FosB staining, slices were incubated in biotinylated secondary antibody (Biotinylated Goat anti-Rb 1:400 in blocking solution; Vector Laboratories, BA-1000) for 1 h. Sections were washed (5×5 min) in PBS then incubated in avidin–biotin complex using ABC Vectastain kit, diluted 1:800 for 1 h. Following washes, sections were incubated in diaminobenzadine (DAB, Pierce, Rockford, IL) and H_2_O_2_ for approximately 10 min, then washed (5×5 min) in saline-free PB. Sections were mounted onto Superfrost Plus microscope slides (Fisher #12-550-15) followed by dehydration in Xylene solution, then slides were cover-slipped using DPX (Sigma, 44581). For Iba1 immunofluorescence staining, sections were incubated in Cy3 secondary antibody (Cy3 anti-Rb 1:500 Jackson, #711-165-152) for 1 h, then washed (5×5 min) in saline-free PB, mounted, and cover-slipped using Gelvatol (Sigma-Aldrich #10981-100 ml). To maintain consistency between samples for each stain, all experimental sections were processed for immunostaining in the same run at the same time.

#### 8.1 Imaging, Quantification and Analysis

Mounted sections were imaged using AxioImager ZI microscope (Zeiss) with apotome (z-stack) capability (Axiocam MRM camera and AxioVision Release 4.6 software; Zeiss). Fear regulatory brain areas were analyzed (infralimbic, IL; prelimbic, PL; basolateral amygdala, BLA; central nucleus amygdala, CeA; dentate gyrus, DG; basal nucleus of the stria terminalis, BNST). FosB+ cells were quantified as previously described via ImageJ

### 9.0 RNA-scope (in situ hybridization)

To fluorescently label RNA, in-situ hybridization was employed using the ACD RNAscope Multiplex Fluorescent v2 kit. Mice were perfused with cold, sterile 1x PBS pH 7.4 (Gibco), and the brains were removed and flash frozen with isopentane. Brains were kept at −80°C until sectioned on a Microm HM500M Cryostat at a thickness of 18µm. Slices containing the SFO and IL were directly mounted onto Superfrost Plus glass slides (Fisher) and stored at −80°C. RNAscope hybridization steps were carried out following the manufacturer’s instructions for fresh frozen tissue. Probes for the *IL-17Ra* (403741, ACD Biosciences), *Iba-1* (Aif1) (319141-C2, ACD Biosciences), and *Gfap* (313211-C3, ACD Biosciences) were used to label IL-17 receptor A, myeloid cells (predominantly microglia in the SFO^24^), and astrocytes, respectively. Immunohistochemistry was performed for neurons using anti-NeuN antibody (ABN78, Sigma-Aldrich) after the RNAscope hybridization as described above to identify co-expression of *IL-17Ra* with SFO neurons. Sections were stained with DAPI (4′,6-diamidino-2-phenylindole) for cell analysis. Images were obtained for RNAscope/IHC samples on a AxioImager ZI microscope (Zeiss) and *Z*-stacks were merged using ImageJ for analysis and quantification of co-localized puncta. Puncta for each probe were hand counted for all cells within the SFO and IL. Cells containing 5 or more puncta for the Iba-1 or Gfap probe were identified as microglia or astrocytes, respectively. When 5 or more IL-17Ra puncta were colocalized with the Iba-1 or Gfap puncta, the cell was considered positive for IL-17Ra. In addition, cells stained positive for the neuronal marker NeuN were counted and analyzed for colocalization with the IL-17Ra probe.

### 10.0 RNAseq

SFO punches were collected from mild and severe AI mice for bulk RNA sequencing at 3 h post HDM. This earlier timepoint was chosen to assess early upstream transcripts and signaling molecules that may drive allergen-associated immune alterations and behavior. SFO enriched tissue was dissected and snap-frozen on dry ice in RNAase-free microcentrifuge tubes and stored at −80°C until processing. Total RNA from each SFO per animal was isolated using RNeasy Mini Kit (Qiagen, #74104) following manufacturer’s directions including genomic DNA removal. RNA concentration was quantified via Nanodrop to 16ug/ml. Samples were stored at −80C until processing. Sequencing was conducted at the University of Cincinnati Genomics, Epigenomics and Sequencing Core utilizing the standard Illumina pipeline ^42.,43^. All samples achieved a RIN >7. For initial analysis, RNA-seq Alignment v2.0.2 (STAR) was run via SEQUENCE HUB followed by RNA-Seq Differential Expression version 1.0.1 (Salmon) using transcripts per million (TPM). Once the sequencing was completed, fastq files were generated via Illumina BaseSpace Sequence Hub. A standard differential expression analysis was performed using DESeq2. Samples were then ranked by fold change and top 10% up and downregulated genes were selected for Enrichr analysis. Top 10 up and downregulated pathways, as well as leading edge genes were identified as the largest drivers of enrichment score (ES) of a given pathway.

### 11.0 Patch Clamp Electrophysiology

Briefly, mice were anesthetized with isoflurane and rapidly intracardially perfused with ice-cold *N*-methyl-*D*-glucamine (NMDG) solution containing (in mM) 92 NMDG, 2.5 KCl, 1.25 NaH_2_PO_4_, 30 NaHCO_3_, 20 HEPES, 25 Glucose, 5 L-ascorbate, 2 thiourea, 3 sodium pyruvate, 10 MgSO_4_, and 0.5 CaCl_2_. 300 µM coronal sections were cut on a VF-300-0Z Compresstome (Precisionary Instruments) into ice-cold NMDG solution. Sections were immediately transferred to room temperature (22-24°C) NMDG solution to recover for 30 min then transferred for a long-term recovery in RT artificial cerebrospinal fluid (aCSF; in mM) 92 NaCl, 2.5 KCl, 1.25 NaH_2_PO_4_, 30 NaHCO_3_, 20 HEPES, 25 Glucose, 5 L-ascorbate, 2 thiourea, 3 sodium pyruvate, 10 MgSO_4_.7H_2_O, and 0.5 CaCl_2_.2H_2_O. Slices containing the subfornical organ (SFO) were transferred to a recording chamber and visualized on an upright microscope (Olympus BX51WI) where they were continually perfused with room temperature extracellular recording fluid containing (in mM) 124 NaCl, 2.5 KCl, 1.2 NaH_2_PO_4_, 24 NaHCO_3_, 5 HEPES, 12.5 Glucose, 2 MgSO_4_, and 2 CaCl_2_, adjusted to pH 7.38 with NaOH and 310 mOsm with sucrose. Recovery chambers and extracellular recording fluid were constantly bubbled with 95% O_2_, 5% CO_2_.

Whole cell patch-clamp recordings from neurons in the SFO were carried out as previously described ^44^, using a Multiclamp 700B patch-clamp amplifier and Digidata 1550 acquisition system with pCLAMP 10 software (Axon instruments). Recordings were sampled at 20 kHz and filtered at 10 kHz. Recording electrodes (4-6 MΩ) were pulled from borosilicate glass capillaries and filled with a potassium-based internal solution containing (in mM) 130K-gluconate, 5 KCl, 5 NaCl, 2 MgCl_2_, 0.3 ethylene glycol-bis(2-aminoethylether)-NNN’N’-tetraacetic acid, 10 HEPES, and 2 Na ATP, adjusted to pH 7.3 with KOH and 294 mOsm with sucrose. Following giga-ohm seal and successful whole cell access, neurons were allowed to stabilize for 2 min then resting membrane potential was recorded. Neurons were then current-clamped at −60mV while measuring membrane properties. After baseline experiments, IL-17A (10ng/mL, eBioscience) was bath applied while resting membrane potential was monitored. Dose-dependent effects of IL-17A on activation of cortical neurons have been reported previously^45^ with optimal effects at the chosen dose. Cells were bathed in IL-17A continuously for 3 min before membrane properties were recorded again.

### 12.0 Stereotaxic Surgery and intersectional chemogenetics

Mice were anesthetized with isoflurane and given pre-operative carprofen (20mg/kg s.c.). For chemogenetic manipulation experiment mice were bilaterally injected with 200nl of Cre-expressing retrogradely transported viral construct - pENN.AAV.hSyn.HI.eGFP-Cre.WPRE.SV40 (Addgene #105540-AAVrg) into the infralimbic cortex (coordinates: A/P 1.54, M/L 0.5, D/V −2.8) to enable Cre expression in IL projecting neuronal soma. For Cre-dependent expression of DREADDs in SFO neurons projecting to the IL, 50 nl of pAAV-hSyn-DIO-hM3D(Gi)-mCherry (Addgene #44362-AAV2) was delivered into the SFO (A/P 0.2, M/L 0.5, angled 80, D/V 2.5). Following recovery (4 wks), mild AI (HDM + IgG) or severe AI (HDM + aC5aR1) was induced followed by behavioral testing at 72 h post HDM. For circuit manipulation mice received either saline or DREADD ligand clozapine oxide, CNO, i.p, at the 2^nd^ and 3^rd^ HDM treatment timepoints, as well as, 24h and 48h post HDM. FC-EXT testing was conducted at 72h.

### 13. Statistical Analyses

Data analysis was conducted using GraphPad Prism 8.2.1 (La Jolla, California) and expressed as mean ± SEM. Outliers were tested via ROUT analysis and excluded from the analysis. One- or two-way ANOVA was conducted with repeated measure ANOVA (RM-ANOVA) analysis applied when applicable. Following a significant main effect, posthoc comparisons were made using the two-stage linear step-up procedure of Benjamini, Krieger and Yekutieli’s with false discovery rate (FDR) correction analysis. RTqPCR and FosB, data were tested for normality via Shapiro-Wilk normality test and standard deviations were compared using the Brown-Forsythe test. Data that passed normality and equal variances were analyzed via ANOVA or t-test where applicable. Graphs were constructed in GraphPad Prism and portions of figures were generated using Biorender.

## Results

### Impaired extinction of fear in mice with allergen-induced severe but not mild airway inflammation: reduced neuronal activation in the infralimbic cortex of severe AI mice

Mild allergen-induced airway inflammation was induced by intratracheal administration of the common aeroallergen house dust mite (HDM), whereas more severe inflammation was induced following combined HDM and antibody-mediated complement C5a receptor antagonism (αC5aR1) (see Fig 1A,B). As expected, compared to PBS-exposed mice (PBS + IgG), those treated with HDM and Isotype control antibody (HDM + IgG) developed airway hyperresponsiveness (AHR) (Fig 1C) and airway eosinophilia (Fig 1D), while limited recruitment of neutrophils, monocytes or lymphocytes was observed (Fig 1E-G). Compared to the HDM + IgG group, mice treated with HDM + αC5aR1 displayed significantly elevated levels of AHR (Fig 1C), and increased recruitment of neutrophils (Fig 1E), monocytes (Fig 1F) and lymphocytes (Fig 1G) with no difference in eosinophil recruitment (Fig 1D). These data suggest that antagonism of C5aR signaling during allergen exposure results in more severe allergen-induced airway inflammation, consistent with previous observations^33^. To assess the immune cell types recruited to the airways and brain, as well as the cytokines produced by these cells in HDM-IgG versus HDM + αC5aR1 groups, we performed flow cytometric analysis on cells isolated from the lungs and whole brain (Figs S1-4). Importantly, there were no differences in the frequency, or number, of IL-13-producing CD4+ T cells or innate lymphoid cells (ILCs) in the lung when HDM + IgG and HDM + αC5aR1-treated mice were compared (Fig S2C,D). While the frequency and number of IL-17A-producing CD4+ T cells was comparable between groups (Suppl Fig 2A,B), a significantly increased frequency (Fig S2E) and total number (Fig S2F) of IL-17A-producing ILCs was observed in the HDM + αC5aR1 severe AI group. There were no significant differences detected in the frequency of immune cell types recruited to the brain (Fig S4). Collectively, our observations of elevated AHR, increased granulocytic inflammation and frequency of IL-17A-producing ILCs, demonstrate that C5aR antagonism in Balb/c mice simulates a severe model of HDM-induced airway inflammation associated with increased IL-17A production in the airways. We henceforth refer to HDM + IgG and HDM + αC5aR1 groups as “mild” and “severe” airway inflammation (AI) respectively.

Behavioral assessments were undertaken in mice with mild versus severe AI compared to control groups: PBS+IgG (CTRL), PBS+αC5aR1 (Ab-CTRL) for effects on fear conditioning-extinction, FC-EXT (Fig 1H), as well as defensive behaviors relevant to anxiety, depression and panic (Fig S5). For FC-EXT, no significant group differences were noted for acquisition (Fig 1I), suggesting that mild or severe airway inflammation does not affect fear learning. Re-exposure to the shock context for conditioned fear (CF) (Fig 1I) also showed no difference in freezing between groups suggesting that associative learning of contextual fear was not impacted by mild or severe AI. To assess extinction learning (E1) and retrieval (E5) mice were exposed to the context for the next 5 days. As evident in Fig 1I, severe AI mice showed higher extinction freezing compared to all other groups while mild AI mice elicited comparable freezing to the control groups. Significantly higher freezing in severe AI mice was observed during early extinction learning (E1) (Fig 1J) and late extinction retrieval (E5) (Fig 1K) compared to all other groups while mild AI mice showed no significant difference from both control groups highlighting that severe AI compromises the extinction of fear. We further assessed the distribution of mice within groups that achieved optimal extinction (defined as <10% freezing at retrieval) (Fig 1L). Notably, ≥50% of mice in the control and mild AI groups achieved extinction (CTRL = 54.5%; Ab-CTRL = 66.6%, Mild = 50.6%;), however only 25.0% of severe AI mice achieved successful extinction. Collectively, these data show that severe (but not mild) airway inflammation compromises fear extinction but maintains the learning and consolidation of fear. To determine contributory brain areas that may regulate observed extinction effects in severe AI mice, we performed post extinction ΔFosB, a proxy for sustained neuronal activation in mild versus severe AI mice. Consistent with the compromised extinction phenotype, a significant reduction in FosB cell counts was observed in severe AI mice within the infralimbic cortex a key extinction regulatory area (Fig 1M), while no significant differences were noted in other fear regulatory areas such as the prelimbic cortex, basolateral amygdala, central nucleus of amygdala, bed nucleus of stria terminalis and hippocampal dentate gyrus (Fig S6). These data support an involvement of the IL cortex in severe AI-induced extinction deficits.

To determine whether the effects of severe AI generalize to other threats, in a separate cohort we tested defensive behaviors to exteroceptive (elevated zero maze, EZM, forced swim test, FST) and interoceptive (CO_2_ inhalation) threat exposures. (Fig S5). No significant group differences were observed in the EZM open arm time Fig S5B). or the FST (Fig S5C) percent immobility or the latency to immobility. Exposure to CO_2_ inhalation-context conditioning paradigm (Fig S5D) revealed significantly increased freezing during CO_2_ inhalation and 24h post CO_2_ context exposure in all groups relative to habituation freezing (Fig S5E), however no significant treatment group differences were noted during habituation, CO_2_ or -context re-exposure days. Collectively these data suggest that severe AI does not impact behaviors related to anxiety-relevant approach-avoidance, learned helplessness, or interoceptive fear and that effects on fear extinction may be selective.

### Necessity of IL-17A for fear extinction retrieval deficit in severe AI mice

Since IL-17A is associated with the severity of airway inflammation ^32,33^ and increased IL-17A secreting cells were observed in severe AI mice, we examined whether IL-17A was associated with compromised extinction in severe AI mice. Since the presence of IL-17A during development has behavioral consequences ^46,47^ and effects of global IL-17A deficiency on brain function are not known, we opted for IL-17A antagonism during allergen administration. Anti-IL-17A antibody (αIL-17A) or IgG was administered to severe AI mice followed by FC-EXT (Fig 2A). No significant group difference was observed for fear acquisition (Fig 2B) or contextual fear memory, CF 24 h later (Fig 2B) suggesting that IL-17A blockade did not impact fear learning or contextual fear memory. αIL-17A treated severe AI mice showed reduced freezing during extinction (Fig 2B). Analysis of early extinction learning (E1) (Fig 2C) revealed no significant group difference, however, severe AI mice treated with αIL-17A showed significantly reduced freezing at extinction retrieval (E5) (Fig 2D). Consistent with improved extinction in αIL-17A treated mice, a distribution analysis for achieved extinction revealed that a large majority of αIL-17A treated severe AI mice successfully achieved extinction (80.0%) while only 36.4% of IgG treated severe AI mice achieved successful extinction. Collectively, these data support that severe AI drives deficits in extinction retrieval via IL17A-associated mechanisms.

**Figure 2:**
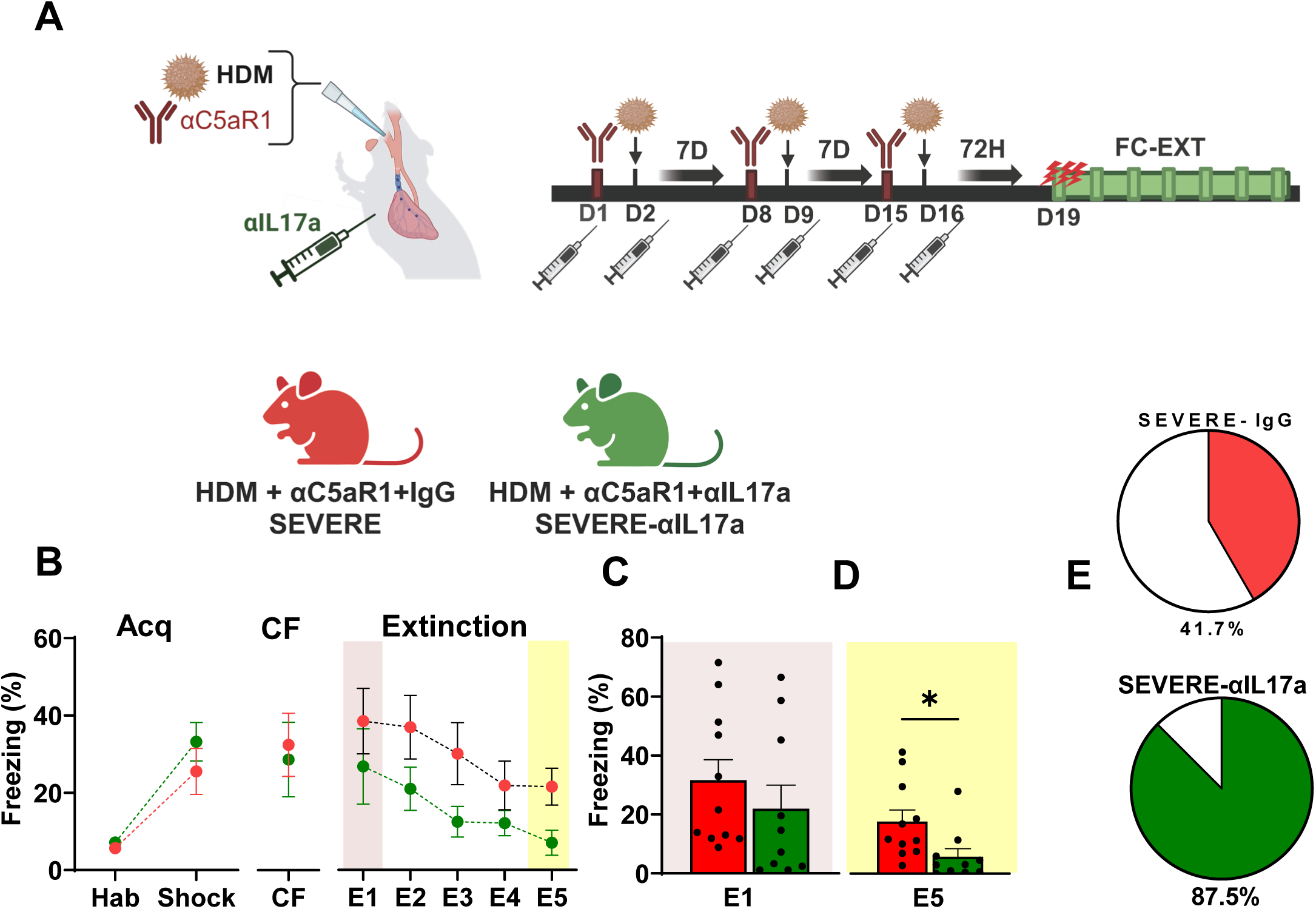
Necessity of IL-17A for fear extinction retrieval deficit in severe AI mice. (A) Mice were given intratracheal administration of PBS/HDM +/− IgG or αC5aR1 intratracheally. Groups received either IgG or anti-IL-17A antibody (αIL17A) intraperitoneally. Schematic illustrates the experimental layout and timeline for treatment and FC-EXT behavior. Experimental groups: HDM-αC5aR1-IgG (severe-IgG) and HDM-αC5aR1-αIL-17A (severe-αIL-17A) (B) For fear acquisition no significant group difference was observed in freezing between the severe-IgG and severe-αIL-17A mice, RM-ANOVA time [significant effect of time, (F_1,38_ = 34.51 p<0.0001) but no effect of treatment (F_1,38_ = 1.369; p = 0.2494) or interaction (F_1,38_ = 0.6317; p = 0.4317). No group differences were noted for contextual fear memory (CF) 24 h later (t_19_ = 0.3010; p = 0.7667). For extinction, αIL-17A-treated severe AI mice showed reduced freezing. Fear extinction over 5 days [time (F_2.094, 39.78_ = 7.645; p = 0.0013), treatment (F_1,19_ = 3.065; p = 0.0961), time x treatment interaction (F _4, 76_ = 0.3179; p=0.86). (C) Freezing during early extinction learning (E1) was not significantly different between severe IgG and severe-αIL-17A groups (t_19_ = 0.9113; p = 0.3735). (D) Severe AI mice treated with αIL-17A showed significantly lower freezing during late extinction retrieval (E5) (t_19_ = 2.453; p = 0.024). (E) Pie charts showing the percent distribution of mice within each group that achieved extinction (defined as <10% freezing on E5). Data are represented as mean ± sem. N= 11 (severe -IgG), 10 (severe-αIL-17A) mice. (*p<0.05)

### Severe AI mice elicit significant microglial activation and shifts in immune transcriptomic signatures within the sensory circumventricular organ, subfornical organ (SFO)

To explore potential brain nodes that may participate in severe AI effects on fear extinction, we assessed microglia, CNS-resident innate immune cells that can sense and transduce immune signals into behavior. CVOs such as the SFO are considered as gateways for body-to-brain communication and previous findings from our lab reported a role of microglia within the circumventricular organ, SFO, in panic-relevant fear^48^. Microglial morphological assessments were undertaken using IBA-1 within sensory circumventricular organs (CVOs), subfornical organ, organum vasculosum laminae terminalis (OVLT) and area postrema (AP), as well as the IL cortex, in mice with severe and mild AI in comparison with CTRL and Ab-CTRL groups at the 72h post allergen timepoint (Fig 3A). Microglial morphological alterations (significantly increased soma perimeter) was observed in the SFO of severe AI mice (Fig 3B, C). compared to mild AI and control groups. No significant microglial alterations were observed within the OVLT, AP or the IL (Fig 3B,C). These data suggest that the SFO is unique among the sensory CVOs in responding to severe AI induced inflammatory response. To determine transcriptional signatures within the SFO that may drive SFO microglial activation in severe AI mice, we conducted bulk-RNAseq in SFO punches comparing severe versus mild AI cohorts (Fig 4A). A volcano plot representation of differentially expressed genes (DEGs) (Fig 4B) revealed significant alterations in several regulatory genes associated with immune signaling and microglial activation in severe AI mice. Transcripts such as Notch1 and interferon regulatory factor 4 (Irf5) that regulate the differentiation and polarization of microglia ^49,,50^ were significantly altered. Ido1, an enzyme linked with microglia phagocytic activity and control of inflammatory response^51^ was upregulated in severe AI. Interestingly, transcripts related to the NK-κ signaling pathway (nuclear factor kappa light chain enhancer of activated B inhibitor beta (*Nfkbib*) and *Rela*) were upregulated in severe AI mice. Notably, IL-17A signaling robustly activates the NF-κB pathway following engagement of its receptor^52^. Several other genes associated with innate immune activation (*S100a10*), endothelial barrier function (*Lama5, Eln, Mmp28, Cldn14*) and cellular trafficking (*Itgb4, S1pr4, Cxcl5*) also showed differential expression between mild and severe AI groups. Furthermore, targeted pathway analysis (EnrichR) revealed 66 differentially altered pathways (23 up- and 43-down-regulated) in the SFO of severe AI mice. Top 10 up- and down regulated pathways (Fig 4 C,D) revealed several pathways associated with immune signaling and responses. Collectively, these data highlight an engagement of the SFO and SFO microglia in severe AI immune responses

**Figure 3:**
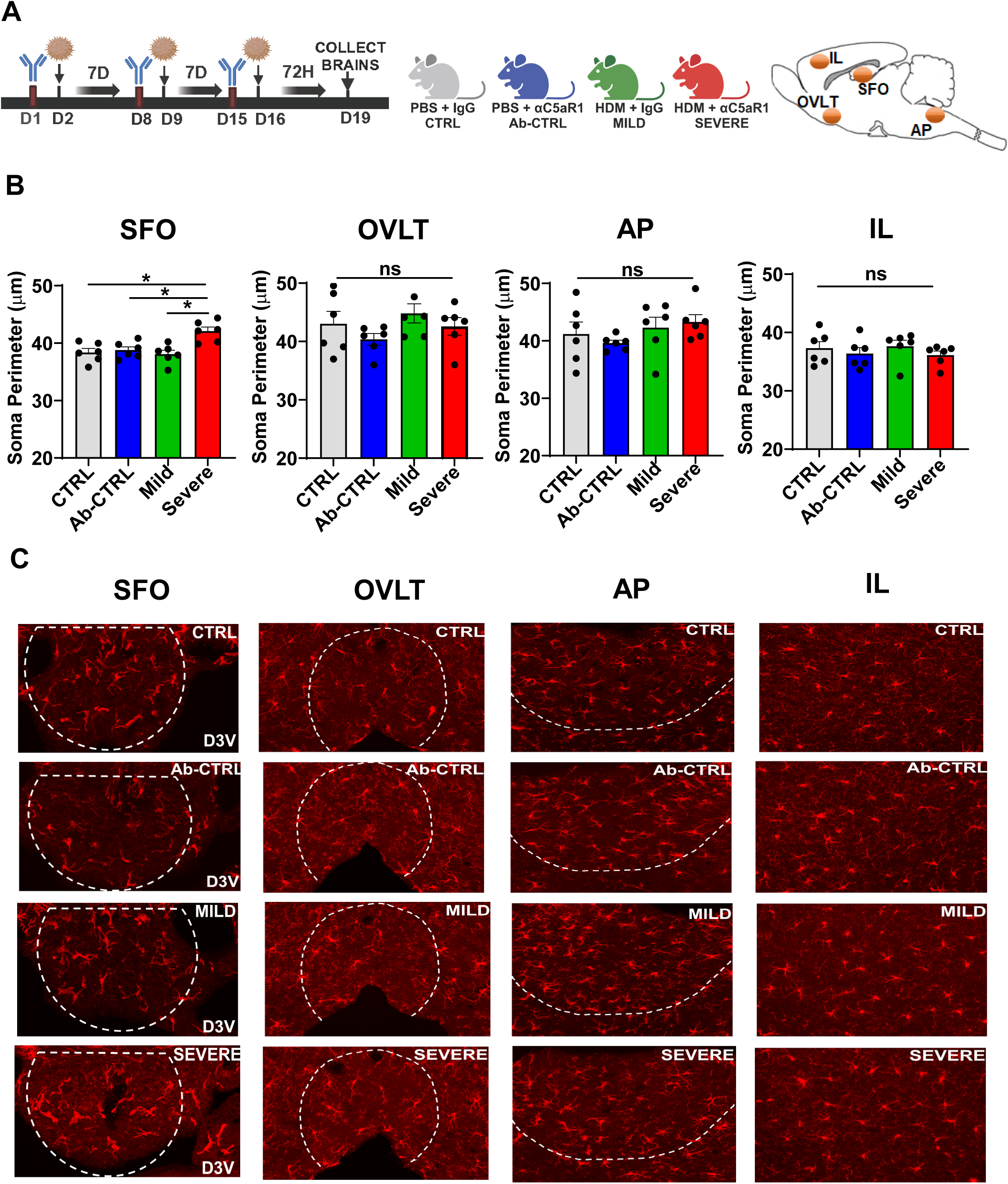
Severe AI mice elicit significant microglial activation within the sensory circumventricular organ, subfornical organ (SFO) (A) Schematic showing the experimental layout and timeline. Mice were intratracheally administered HDM ± IgG or αC5aR1 to generate mild versus severe AI and compared with control groups receiving PBS-IgG (CTRL) or PBS-αC5aR1 (Ab-CTRL). Mice were perfused and brain isolated for immunohistochemical staining for IBA-1 in sensory CVOs: subfornical organ (SFO), organum vasculosum laminae terminalis (OVLT) and area postrema (AP), as well as the infralimbic (IL) cortex. (B) Morphological alterations were observed in microglia within the SFO of severe AI mice (1-way ANOVA: F_3,20_ = 7.601, p=0.0014). Post hoc comparisons revealed significantly increased soma perimeter in severe AI mice compared to mild AI and control groups (severe vs CTRL p=0.0045; severe vs Ab-CTRL p= 0.012; severe vs mild p= 0.002). No significant morphological alterations in microglial soma perimeter were observed within the OVLT (F_3,20_ = 1.286, p=0.360), AP (F_3,20_ = 1.014; p=0.405) or the IL (F_3,20_ = 0.7314; p=0.545) (C) Representative images of IBA-1 positive cells within quantified brain areas: SFO, OVLT, AP and IL from CTRL, Ab-CTRL, mild AI and severe-AI mice, D3V= dorsal third ventricle Data are represented as mean ± sem. N= 6 mice per group (*p<0.05)

**Figure 4:**
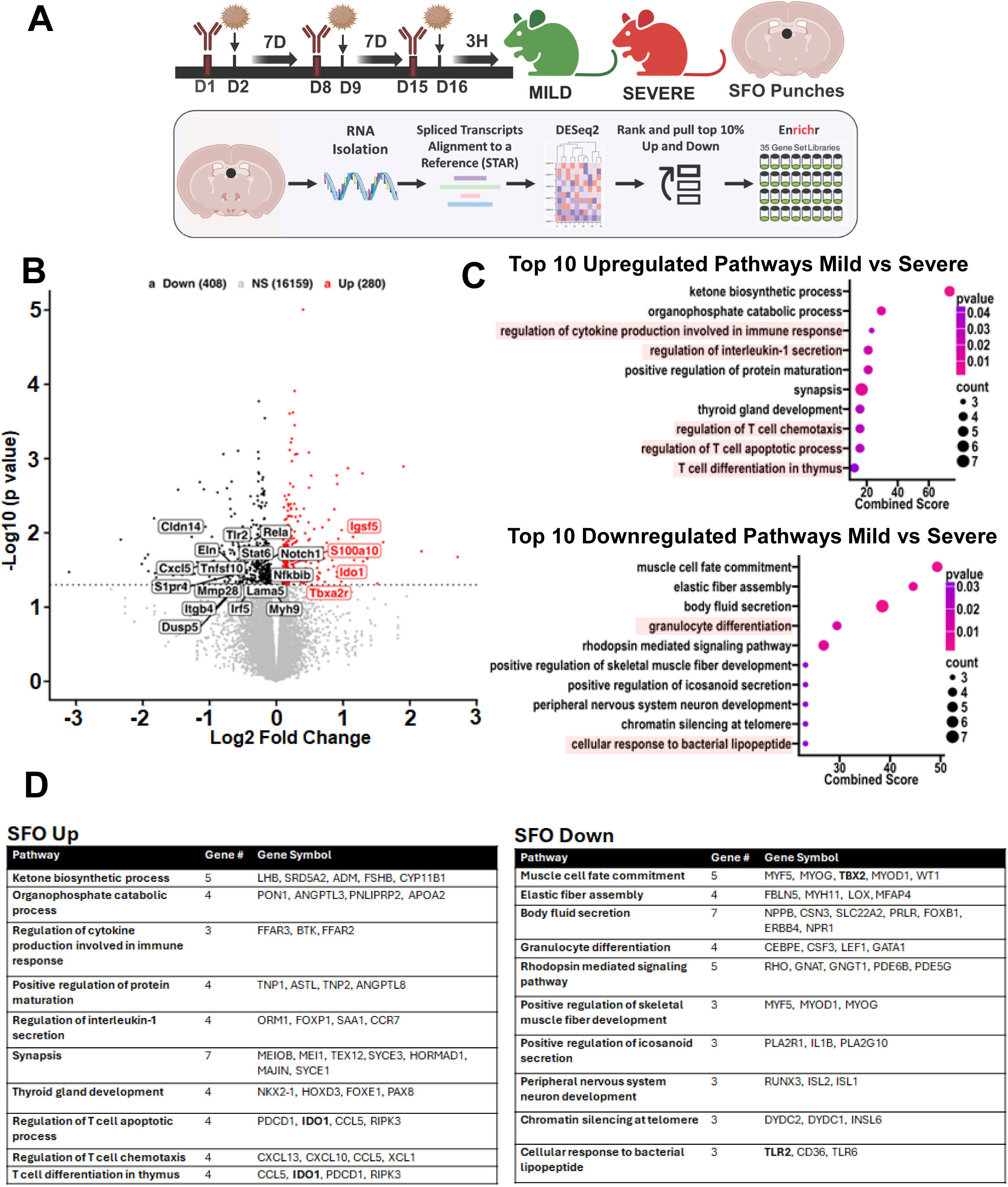
Severe AI mice elicit shifts in immune transcriptomic signatures within the subfornical organ (SFO) compared to mild AI mice. (A) Schematic for experimental layout. Mice were intratracheally administered HDM-IgG or HDM-αC5aR1 to generate mild versus severe AI, respectively. To determine transcriptional signatures that may drive downstream effects, SFO enriched tissue punches were collected at 3h following the last allergen administration. Bottom panel shows sample processing for sequencing and analysis to yield top 10% up-and down-regulated genes that underwent Enrichr analysis for the identification of top 10 up- and down-regulated pathways. (B) Volcano plot representation of differentially expressed genes (DEGs). Immune genes differentially expressed in severe AI mice included genes associated with immune signaling and microglial activation (Irf5, Notch1, Ido1, Nfkbib, Rela, S100a10), endothelial cell barrier function (Lama5, Eln, Mmp28, Cldn14) and cellular trafficking (Itgb4, S1pr4, Cxcl5). (C) Pathways identified via Enrichr analysis revealed 66 differentially altered pathways (23 up- and 43-down-regulated) in the SFO of severe AI mice. Top 10 up- and down regulated pathways are shown here and included several pathways associated with immune signaling and response (see highlighted) (D) Tables show leading edge genes associated with top 10 up- and down-regulated pathways.

### SFO microglia express IL-17 receptor A (IL-17RA): Microglial Il17ra deficient severe AI mice have improved fear extinction

Previous studies have highlighted a regulatory role of the brain IL-17 receptor A subunit, IL-17RA in neuroinflammatory models^53,54^ and neuronal IL-17RA regulates anxiety^45^ and social^46,47^ behaviors. Currently, functional contributions of microglial IL-17RA are not well understood and its role in behavioral regulation has not been studied. Given the association of IL-17A with compromised extinction and an engagement of the SFO in severe AI mice, we investigated the cellular expression profile of the IL-17RA within the SFO. Using a high throughput droplet-based-single-cell RNA sequencing dataset reported previously^55^, we mapped the expression of the IL-17RA transcript on SFO cell type clusters. An unsupervised clustering on graph-based representation of cellular gene expression profiles visualized in a uniform manifold approximation and projection (UMAP) embedding revealed diverse cell types characterized by unique transcriptional signatures (t-SNE plot (Fig 5A). A feature plot of cell-type specific expression of *Il17ra* (Fig 5B) revealed a glia-skewed expression of *Il17ra* localized to microglia as well as LT astrocytes. Additionally, we identified two microglial clusters, microglia-1 and microglia-2 in the SFO with distinct transcriptional signatures (Fig S7A,B). Violin plot analysis revealed enriched expression of IL-17RA in microglia-1 cluster (Fig 5C). Interestingly, while canonical microglial genes were expressed in both populations, microglia-1 cluster showed enriched expression of genes associated with phagocytosis, immune activation, cytokine/chemokine signaling, TLR/inflammasome signaling relative to microglia-2 (see Fig S7) suggesting that this cluster may represent SFO microglia that are more receptive to immune signals.

**Figure 5:**
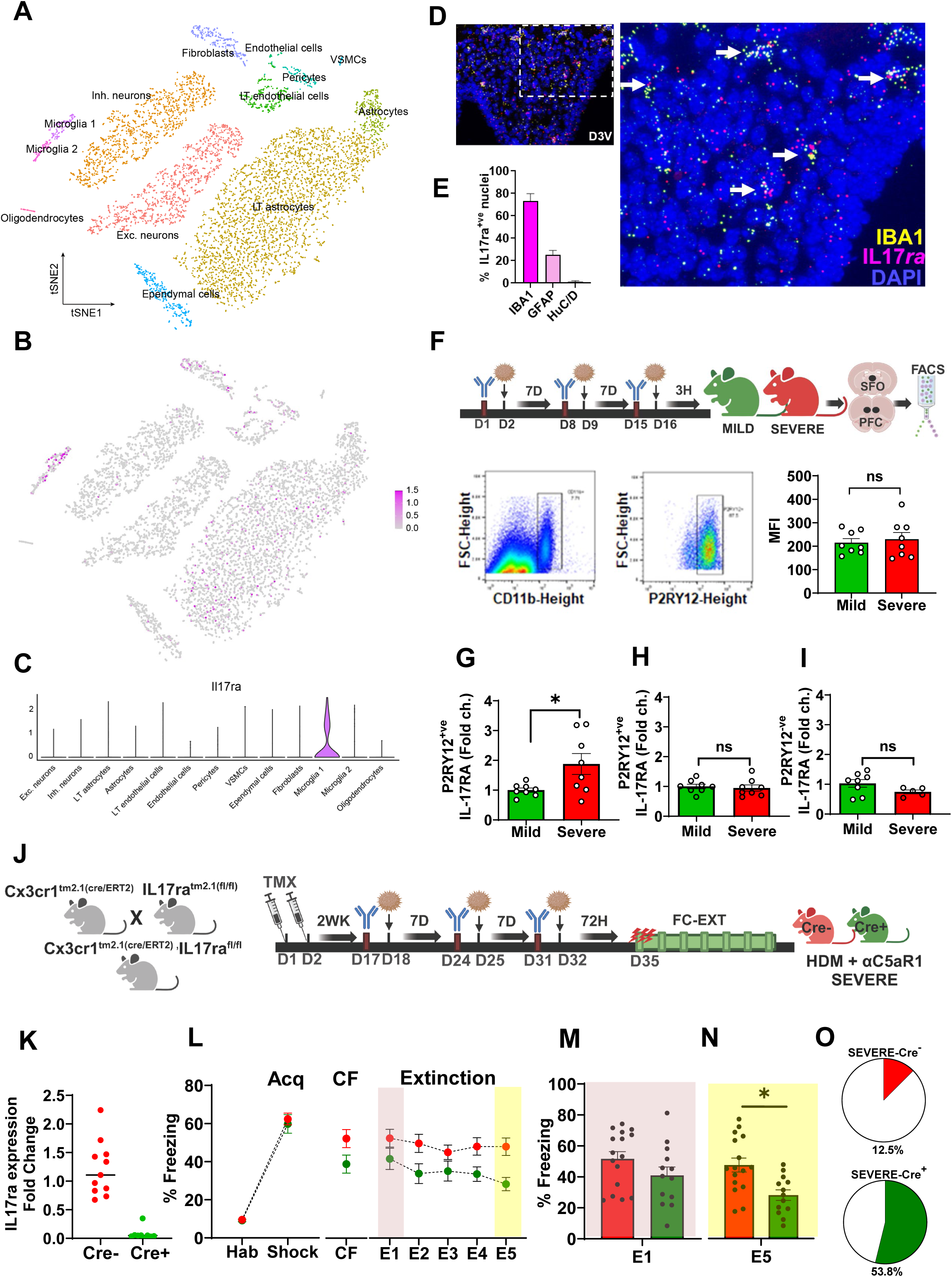
SFO microglia express IL-17A receptor (IL-17RA): Microglial *Il17ra* deficient severe AI mice have improved fear extinction. (A) Single-cell RNA-sequencing analysis of cellular diversity in the SFO reveals 13 transcriptomic cell classes with two microglial populations. Data shown in a tSNE embedding of 7,950 cells with color-coded cell identity. (B) Log-normalized expression of Il17ra in SFO cell classes. (C) Violin plot of log-normalized expression of Il17ra in SFO cell classes. Data in A-C reanalyzed and plotted from a previous study (Pool et al, 2020) [56]. (D) Representative image from RNAscope *in-situ* hybridization on SFO slices using probes for *Il17ra* and ionized calcium binding protein *IBA1* showing co-localization of *IL17ra*^+ve^ puncta (magenta) with IBA1^+ve^ puncta (yellow) and DAPI stained cells (blue). Inset shows low magnification 20X image of the whole SFO with the dotted line indicating area shown in the magnified 40X image. D3V= dorsal 3^rd^ ventricle (E) Bar graph shows quantified % colocalization of *Il17ra*^+ve^ puncta with IBA1^+ve^ (73%), GFAP^+ve^ (25.3%) puncta and HuC/D^+ve^ soma (1.1%). Fig S8 shows Images for *Il17ra*-GFAP and *Il17ra*-HuC/D in-situ hybridization. N= 2-5 SFO slices (F) Schematic showing the experimental layout for FACS measurements on SFO and PFC enriched tissue. Mice were intratracheally administered HDM ± IgG or αC5aR1 to generate mild versus severe AI. SFO and PFC tissue were collected 3h following the last HDM administration. Bottom panel shows representative image for gating of CD11b^+ve^ /P2RY12^+ve^ cells. Mean fluorescence intensity (MFI) of samples from mild and severe AI mice was not significantly different. (G) Quantitative PCR (qPCR) assessment of *Il17ra* mRNA expression in FACS sorted P2RY12^+ve^ SFO microglia from severe AI and mild AI mice. Significantly higher *Il17ra* mRNA expression was observed in P2RY12^+ve^ SFO microglia from severe compared to mild AI mice (t test: t_14_ = 2.445, p= 0.028, N= 8 mice per group). *Il17ra* expression was not significantly different in P2RY12^+ve^ microglia from the prefrontal cortex (H) or the P2RY^−ve^ cells from the SFO (I) (J) Strategy for the generation of Cx3cr1^cre/ERT2^: IL17ra^fl/fl^ mice and the experimental timeline showing administration of tamoxifen, followed by a 2-week recovery period to enable establishment of peripheral Cx3cr1^+ve^ macrophages. Subsequently, mice underwent intratracheal HDM ± IgG or αC5aR1 administration followed by behavioral testing for FC-EXT. Control mice were *Il17ra*^fl/fl^ but lacked Cre/ERT2 (Cre^−ve^). (K) FACS sorted P2RY12^+ve^ SFO microglia from Cre^+ve^ and Cre^−ve^ Cx3cr1^cre/ERT2^ *Il17ra*^fl/fl^ mice were analyzed for *Il17ra* mRNA expression using qPCR. Bar graph shows negligible *Il17ra* transcript levels in Cre^+ve^ vs Cre^−ve^ mice N= 11 (Cre^−ve^), 9 (Cre^+ve^). (L) Severe-AI phenotype was generated in Cre^+ve^ and Cre^−ve^ Cx3cr1^cre/ERT2^:*Il17ra*^fl/fl^mice followed by FC-EXT. For fear acquisition, no significant group difference was observed in freezing between the severe-Cre^+ve^ and severe-Cre^−ve^ mice (RM-ANOVA, time (F_1,27_ = 345.7, p<0.0001) but no genotype (F_1,27_ = 0.264; p = 0.611) or interaction (F_1,27_ = 0.154; p =0.69), Conditioned freezing (CF) 24 h later was not significantly different (t_27_ = 1.98; p = 0.06). For extinction, severe-Cre^+ve^ mice showed reduced freezing. Fear extinction over 5 days revealed the effects of time (F_2.242, 60.53_ = 2.614, p=0.07) and treatment (F_1, 27_=6.798, p=0.015) but there was no interaction (F_4,108_= 0.9308, p=0.449) N= 13 (Cre^+ve^), 16 (Cre^−ve^). (M) Freezing during early extinction learning (E1) was not significantly different between severe-Cre^+ve^ and severe-Cre^−ve^ groups (t_27_ = 1.508; p = 0.143). (N) Severe AI-Cre^+ve^ mice showed significantly lower freezing during late extinction retrieval (E5) (t_27_ = 3.28; p = 0.002) as compared to the severe AI-Cre^−ve^ group. (O) Pie charts show the percent distribution of mice within each group that achieved extinction (defined as <30% freezing on E5). Note: In this cohort freezing was higher in FC-EXT, potentially contributed by genotype and strain differences (C57/Bl6). Data are represented as mean ± sem. N= 16 (severe-Cre^−ve^), 13 (severe-Cre^+ve^) mice (*p<0.5)

In-situ hybridization using an *Il17ra* mRNA probe further confirmed microglial *Il17ra* expression in the SFO (Fig 5D, Fig S8A,B). Quantification of *Il17ra* puncta (≥5/cell) (Fig 5E) revealed abundant co-localization with IBA1^+ve^ puncta (73%) compared to GFAP^+ve^ puncta (25.3%) and HuC/D^+ve^ cells (1.1%). In comparison, *Il17ra mRNA* expression within the infralimbic cortex was localized to IBA1^+ve^ cells and HuC/D^+ve^ neurons (Fig S8C-F). Collectively, our single-cell transcriptomics and *in-situ* hybridization analyses indicate enriched microglial *Il17ra* expression in the SFO.

We further examined whether SFO microglial *Il17ra* is regulated in severe AI mice using FACS-sorted microglia. Enriched SFO P2RY12^+ve^ microglia (Fig 5F) revealed a significant upregulation of *Il17ra* in severe AI mice versus mild AI (Fig 5G), an effect that was not observed in P2RY12^+ve^ microglia from the prefrontal cortex (Fig 5H) or P2RY12^−ve^ cells in the SFO (Fig 5I). To test the necessity of microglial IL-17RA in mediating severe AI effects on fear extinction, we generated mice with conditional knockdown of microglial *Il17ra* (Cx3cr1^Cre/ERT2^: *Il17ra*^fl/fl^ mice Fig 5J). Effective Cx3cr1^CreERT2^ mediated deletion of *Il17ra* was observed in FACS-sorted P2RY12^+ve^ SFO microglia (Fig 5K). Following an extended 2-week period post tamoxifen (to enable *Il17ra* expression in peripheral macrophages), Cre^+ve^ and Cre^−ve^ mice underwent HDM/αC5aR1 treatment to generate severe AI, followed by FC-EXT testing (Fig 5J). Severe AI mice with *Il17ra* deficiency elicited no significant differences in fear acquisition and conditioned freezing (CF) 24 h later (Fig 5L). Consistent with our hypothesis, *Il17ra*-Cre^+ve^ severe AI mice demonstrated reduced freezing during extinction (Fig 5L). Analysis of early extinction learning (E1) (Fig 5M) revealed no significant group difference, however (consistent with anti-IL-17A antibody, αIL-17A data) a significant reduction in E5 extinction retrieval freezing was observed in *Il17ra*-Cre^+ve^ mice (Fig. 5N). Furthermore, the distribution of mice with extinction was higher in *Il17ra*-Cre^+ve^ (54%) than *Il17ra*-Cre^−ve^ (12.5%) severe AI mice. Collectively, these data support the recruitment of SFO microglial IL-17RA in regulating severe AI effects on extinction.

### Activation of SFO neurons by IL-17A

IL-17A/IL-17RA-mediated behavioral effects in severe AI mice would require the participation of SFO neurons. We used Fos^Cre/ERT2.Ai9^ mice and patch clamp electrophysiology to determine whether IL-17A can activate SFO neurons. Whole-cell current-clamp recordings from SFO neurons in slice were performed before and following bath application of IL-17A (Fig 6 A-C). Application of IL-17A significantly increased the amplitude of action potentials suggesting overall enhanced excitability (Fig 6C). There was no significant change in the action potential threshold or rheobase. Additionally, SFO-targeted administration of IL-17A in Fos^Cre/ERT2.Ai9^ mice (Fig 6D) led to a significant increase in td-tomato-cFos^+ve^ cells (Fig 6E). Collectively, these data support the activation of SFO neurons downstream of IL-17A.

**Figure 6:**
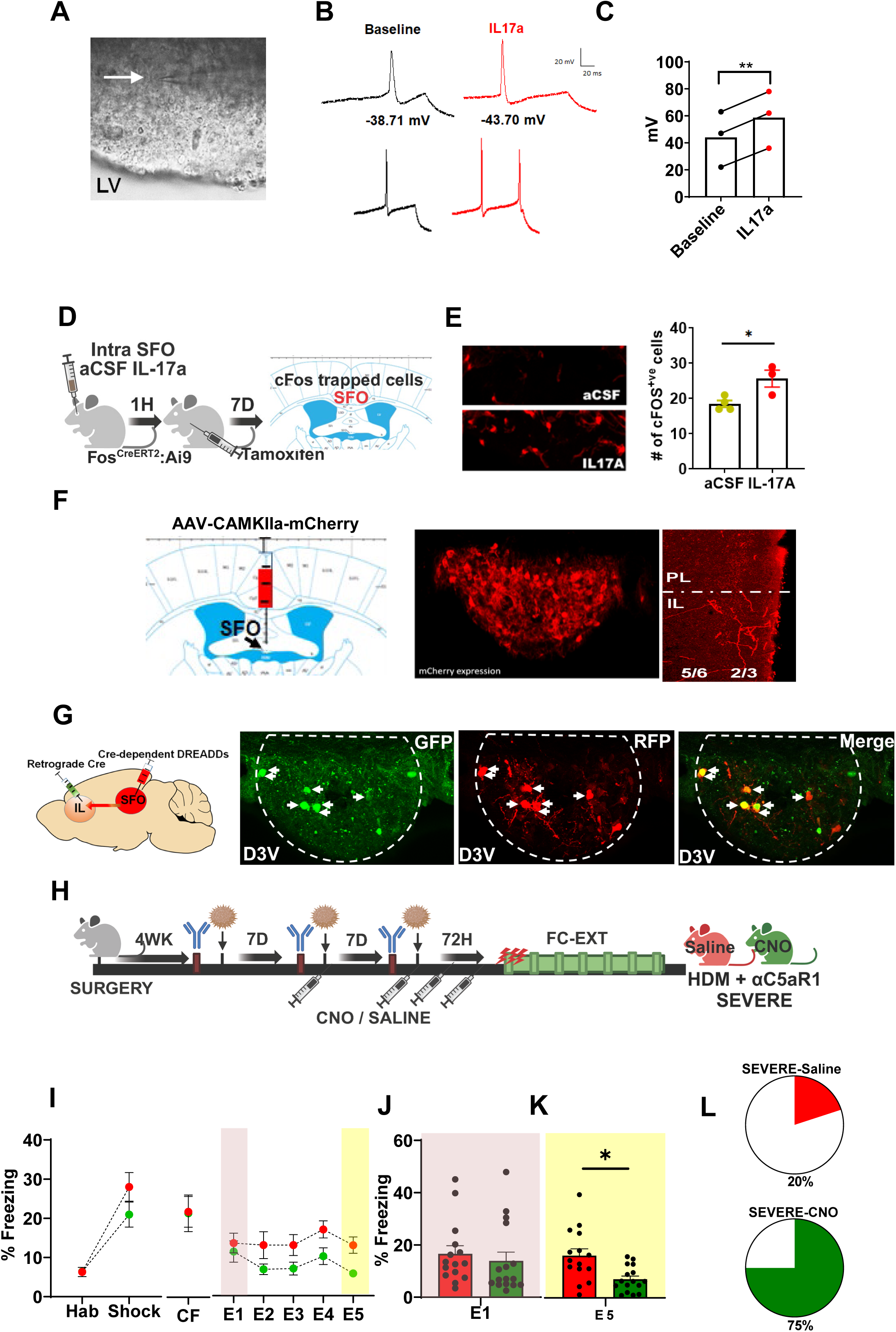
Activation of SFO neurons by IL-17a and SFO to IL circuit regulates fear extinction deficits in severe AI mice. (A) Patch clamp recordings from SFO neurons in current clamp mode at baseline and following an infusion of IL-17A (10 ng/ml). Arrow in panel A shows patched cell with recording electrode; LV = lateral ventricle (B) Recording traces show decreased action potential threshold (top) and increased action potential discharge (bottom) following IL-17A. There was no significant change in the threshold of the first action potential fired, nor was there a change in rheobase. (C) Infusion of IL-17A significantly increased action potential amplitude from baseline (paired t test t_2_ = 44.0; **p= 0.0005 vs baseline) (D) Layout for experiment on intra-SFO infusion of IL-17A or aCSF in Fos ^CreERT2.Ai^^9^ mice. cFos trapped td-tomato^+ve^ cells were visualized in the SFO. (E) Image panels show td-tomato^+ve^ cells in aCSF (top) and IL-17A (bottom) treated mice. Bar graph shows significantly higher # of cFos/td-tomato^+ve^ cells in the IL-17A group (unpaired t test, t_5_ = 3.105; p = 0.026). (F) Visualization of SFO terminals in the medial prefrontal cortex using viral tract tracing. An AAV-CAMKIIa-mCherry construct (Addgene) was delivered into the SFO. After 4 wk, using RFP immunostaining, terminals were observed in the IL outer layers 2/3. No RFP^+ve^ terminals were observed in the adjacent prelimbic (PL) area. (G) Intersectional chemogenetics strategy to target the SFO-IL circuit using a Cre-expressing retrogradely transported viral construct - pENN.AAV.hSyn.HI.eGFP-Cre.WPRE.SV40 into the IL and pAAV-hSyn-DIO-hM3D(Gi)-mCherry virus in the SFO to enable Cre-dependent expression of DREADDs in SFO neurons projecting to the IL. Images show the expression of eGFP in Cre expressing SFO to IL neurons, Cre-dependent RFP DREADD-Gi expressing soma and colocalized cells. D3V= dorsal third ventricle (H) Schematic for experimental setup. Following post-surgery recovery, mice received HDM/αC5aR1 treatments to induce a severe AI phenotype. Mice received either saline or CNO, i.p, at the 2^nd^ and 3^rd^ HDM treatment timepoints, as well as, 24h and 48h post HDM. FC-EXT testing was conducted at 72h. (I) For fear acquisition (Acq), no significant group difference was observed in freezing between saline and CNO-treated severe AI mice (RM-ANOVA: effect of time, (F_1,39_ = 59.91 p<0.0001) but no treatment (F_1,39_ = 1.70; p = 0.20) or treatment x time interaction (F_1,39_ = 2.307; p =0.137)]. Conditioned freezing (CF) 24 h later was not significantly different (t_29_ = 0.064; p = 0.949). For extinction, severe-CNO mice showed reduced freezing. Fear extinction over 5 days revealed the effects of time F _(3.277, 95.04_ = 2.206; p=0.08 and treatment (F _1, 29_= 6.187, p=0.018) but there was no interaction (F _4, 116_= 0.6689, p=0.614) (J) Freezing during early extinction learning (E1) was not significantly different between saline and CNO groups (t_29_ = 0.583; p = 0.564). (K) CNO-treated severe AI mice showed significantly lower freezing during late extinction retrieval (E5) (t_19_ = 3.22; p = 0.003) as compared to the severe AI-saline group. (L) Pie charts show the percent distribution of mice within each group that achieved extinction (defined as <10% freezing on E5). Severe AI-CNO= 75%; severe AI-saline=20%. N= 15 (saline), 16 (CNO) (*p<0.05) All data are mean ± sem, * p<0.05 between indicated groups

### SFO to Infralimbic (IL) cortex circuit regulates fear extinction deficits in severe AI mice

While functional connectivity of the SFO to homeostatic regulatory nodes such as the hypothalamus, medial preoptic nucleus and other CVOs is well established for the regulation of motivated behaviors ^28,56,57^, there is currently no information on the functional role of SFO-cortical circuits. SFO neuronal projection terminals in the prefrontal cortex were visualized using AAV-tract tracing (Fig 6F). In mice with SFO-targeted AAV delivery, terminal fibers were localized in the IL primarily in the outer layers 2/3. No labelling was detected in the adjacent prelimbic area. To determine whether the SFO-to-IL circuit regulates compromised extinction in severe AI mice, an intersectional chemogenetics approach was utilized (Fig 6G,H). Following recovery to enable Gi-DREADD expression in SFO neurons projecting to the IL (Fig 6G), mice received HDM/αC5aR1 treatment and saline or DREADD ligand, CNO, was administered prior to FC-EXT testing (see schematic Fig 6H). No significant group differences were observed during fear acquisition (Fig 6I) or context conditioned freezing (CF) suggesting that the SFO-to-IL circuit manipulation did not impact fear learning or contextual fear memory. Consistent with our αIL-17A (Fig 2) and microglial *Il17ra* ablation data (Fig 5), CNO-treated severe AI mice elicited reduced freezing during extinction (Fig 6I). Analysis of early extinction learning (E1) (Fig 6J) revealed no significant group difference, however, CNO-treated mice froze significantly less during E5 extinction retrieval (Fig 6K). Distribution analysis revealed that a higher number of CNO-treated severe AI mice achieved extinction (75%) compared to saline (20%) (Fig 6L). Collectively, these data strongly support that the SFO-IL circuit regulates fear extinction deficits in mice with severe AI.

## Discussion

Although the association between lung inflammation and poor mental health has long been recognized, the underlying mechanisms remain poorly understood. Here, we demonstrate that severe—but not mild—allergic airway inflammation (AI), induced by house dust mite exposure, impairs the extinction of conditioned fear. We identify an IL-17A–mediated mechanism involving the subfornical organ (SFO), a sensory circumventricular organ (CVO) with direct access to the systemic milieu and highlight a key role for microglial IL-17RA and the SFO-to-infralimbic cortex (IL) circuit in mediating extinction deficits in severe AI.

Chronic airway inflammation, beyond its peripheral consequences, can negatively affect emotional processing and mental health ^2,5,11^. In this study, we focused on allergic asthma–associated AI, the most common form of lung inflammation affecting approximately 300 million individuals worldwide ^58^. Allergic asthma represents about 60% of all asthma cases and is particularly prevalent in children, as approximately 11% of U.S. adolescents aged 15–19 are diagnosed with the condition ^7^, underscoring the impact of our findings.

Our results revealed an association of severe AI with extinction deficits. Severe AI mice elicited pronounced neutrophilic inflammation and increased IL-17A–producing innate lymphoid cells (ILCs). Notably, anti-IL-17A improved extinction in these mice, implicating this cytokine as a key modulator. Prior studies have linked IL-17A to anxiety-like and social behaviors in homeostatic ^45^ and developmental ^46,47^ contexts. Our findings extend its role to extinction memory regulation—underscoring IL-17A as a systemic immune signal capable of shaping emotional behavior.

Interestingly, fear learning and associative memory remained intact in severe AI mice, and responsiveness to other threat paradigms was also unaltered. This selectivity suggests targeted disruption of extinction regulatory mechanisms in severe AI mice, as supported by reduced neuronal activation within the IL, a key area encoding the retrieval of extinguished fear ^59^.

We explored possible afferent mechanisms that may transmit IL-17A effects to the IL for extinction regulation and focused on the CVOs - specialized brain regions that can directly sense circulating signals. Importantly, strategic neural connections to forebrain emotion-regulatory sites^22^ enable CVOs to integrate systemic cues with behavioral regulation. CVOs are reported to detect endotoxins and pro-inflammatory cytokines in modulating sickness and anxiety behaviors ^60–62^. Our data revealed selective activation of innate immune cells, microglia within the SFO (but not OVLT, area postrema or the IL). Altered immune gene signatures in the SFO of severe versus mild AI mice further support its recruitment in severe AI effects.

Since IL-17A was necessary for severe AI-associated extinction deficits we investigated cell types within the SFO that can respond to IL-17A. Neuronal IL-17 receptor A (IL-17RA)-mediated regulation of anxiety and social behaviors has been previously reported under homeostatic^45^ and inflammatory^46,47^ conditions and IL-17a can activate astrocytes and microglia primarily through IL-17RA (but not IL-17RC) signaling ^54,63^. We observed high expression of *Il17ra* in SFO microglia. Notably, *Il17ra* was enriched in a transcriptionally distinct microglial cluster that showed high expression of several genes related to immune activation and signaling. Although cell-type specific *Il17ra* expression was not conducted in other CVOs, presence of *Il17ra* on SFO microglia that are highly receptive to immune signals could explain selective engagement of the SFO in severe AI mice. Recent single-cell transcriptomics data has revealed distinct cellular composition and unique gene expression signatures within the SFO and OVLT ^55^ that may regulate unique functional responses by these areas. Importantly, FACS-sorted microglia from the SFO-but not prefrontal cortex-showed *Il17ra* upregulation in severe AI mice, and microglial *Il17ra* deletion improved extinction, collectively confirming the necessity of this signaling mechanism in severe AI induced extinction deficits. While IL-17RA–dependent microglial activation has been implicated in multiple sclerosis, infection, and neurodegeneration ^64–66^, its role in behavioral regulation is novel. Previously, we also reported acid-sensing by SFO microglia in CO_2_ inhalation evoked fear and panic-like behaviors ^38,39,48^ suggesting a broader chemosensory role for these cells that expands beyond immune signals in regulating emotional behaviors.

Our data also uncovers a previously underappreciated neural pathway from the SFO to the IL that regulates fear extinction. Although CVOs are recognized for their role in interoception and homeostatic regulation ^21,67,68^ their influence on emotional behavior has not been clearly defined. Anatomical studies reported SFO projections to the prefrontal cortex decades ago ^69^, but this is the first demonstration of a functional role for the SFO-IL circuit in modulating fear extinction. Chemogenetic inhibition of this pathway improved extinction retrieval in severe AI mice, indicating that activation of the SFO-IL circuit suppresses IL-mediated fear regulation. Functionally, this circuit may act as an adaptive mechanism that limits cortical emotional processing during systemic inflammatory stress, potentially redirecting neural resources toward physiological homeostasis. This is conceptually parallel to the well-documented suppression of immune responses under extreme psychological stress.

While our findings highlight a novel IL-17A–SFO–IL mechanism, some limitations warrant consideration. Our model focused exclusively on immune-inflammatory mechanisms and did not incorporate the role of psychological stress. Given the SFO’s role in autonomic and neuroendocrine responses^70^, integrating stress response with behavior would provide a more complete understanding. Our study focused on central mechanisms; however, it would be important to establish peripheral IL-17A signaling mechanisms. Additionally, this study included only male mice; considering known sex differences in immune function^71^ and fear regulation^72^, investigating female subjects is a critical next step. We also acknowledge the likelihood of other body-to-brain communication pathways in our model. The vagus nerve provides sensory input from the lungs to the brainstem and is involved in regulating airway reactivity and sickness behaviors ^73,74^. It is plausible that humoral and sensory mechanisms cooperate to coordinate the complex behavioral and physiological responses to inflammation. In this context, the SFO–IL circuit may represent one such conduit specialized for effects on emotional processing.

The implications of our findings are substantial. Chronic, severe airway inflammation is often conceptualized only as a systemic inflammatory disease, yet our data suggest that it may also compromise recovery from trauma, increasing vulnerability to fear-related disorders such as PTSD, phobias, or panic disorder. Our study provides mechanistic insights on the observed co-occurrence of asthma and PTSD^11^, although longitudinal studies are needed to clarify causality. Given the increasing interest in immunopsychiatry^75^, screening PTSD patients for underlying inflammatory conditions—including asthma—and biomarkers like IL-17A may help with treatments and identify at-risk individuals. The SFO’s accessibility to blood-borne signals also presents a unique opportunity for therapeutic targeting to alleviate cortical dysfunction and psychiatric symptoms. Finally, while we used a model of allergic asthma, elevated IL-17A is increased in other pulmonary pathologies linked to poor mental health—such as ARDS and COVID-19—broadening the potential relevance of our findings.

In summary, we identify a novel interoceptive pathway in which severe AI-associated IL-17A signaling in SFO microglia impairs fear extinction via an SFO–IL circuit. This pathway highlights how severe airway inflammation may predispose individuals to fear-related psychopathology, offering new insights into immune-to-brain communication and translational opportunities for treatment.

## Acknowledgments

This work was supported by the National Institute of Health Grants R01 MH133070, R56 MH127043 and R21 MH117483 (RS, IPL - MPIs), R01MH121102 and R01AG083628 (REM), R01 MH123545 and R01 MH130399 (ESW) and NIH R35GM146890 (II). A-H.P was supported by the Eugene McDermott Endowed Scholars Fund. Trainees would like to acknowledge support from NIH T32 training grants: T32NS007453 (EA) and 1T32GM144873-01 (WGR). The authors would like to thank other lab members from our groups: Julie Hargis, Pranav Narayanan, Anisha Ajmani, Maryam Shinwari, Ali Imami and Kristin Oshima for excellent technical assistance.

## Author Contributions

EA, IPL and RS conceptualized the project; EA, JWM, RAA performed majority of the experiments and data analysis with support from EM, LLV, AW, KMJM. Other contributions: LM and SD (SFO electrophysiological recordings), BS and II (brain flow cytometry), WR and REM (SFO bulk RNAseq analysis), A-H P (SFO sc-transcriptomics analysis) and ESW (FACS). EA and RS wrote the paper with input from all other authors.

---

**Fig S1:**
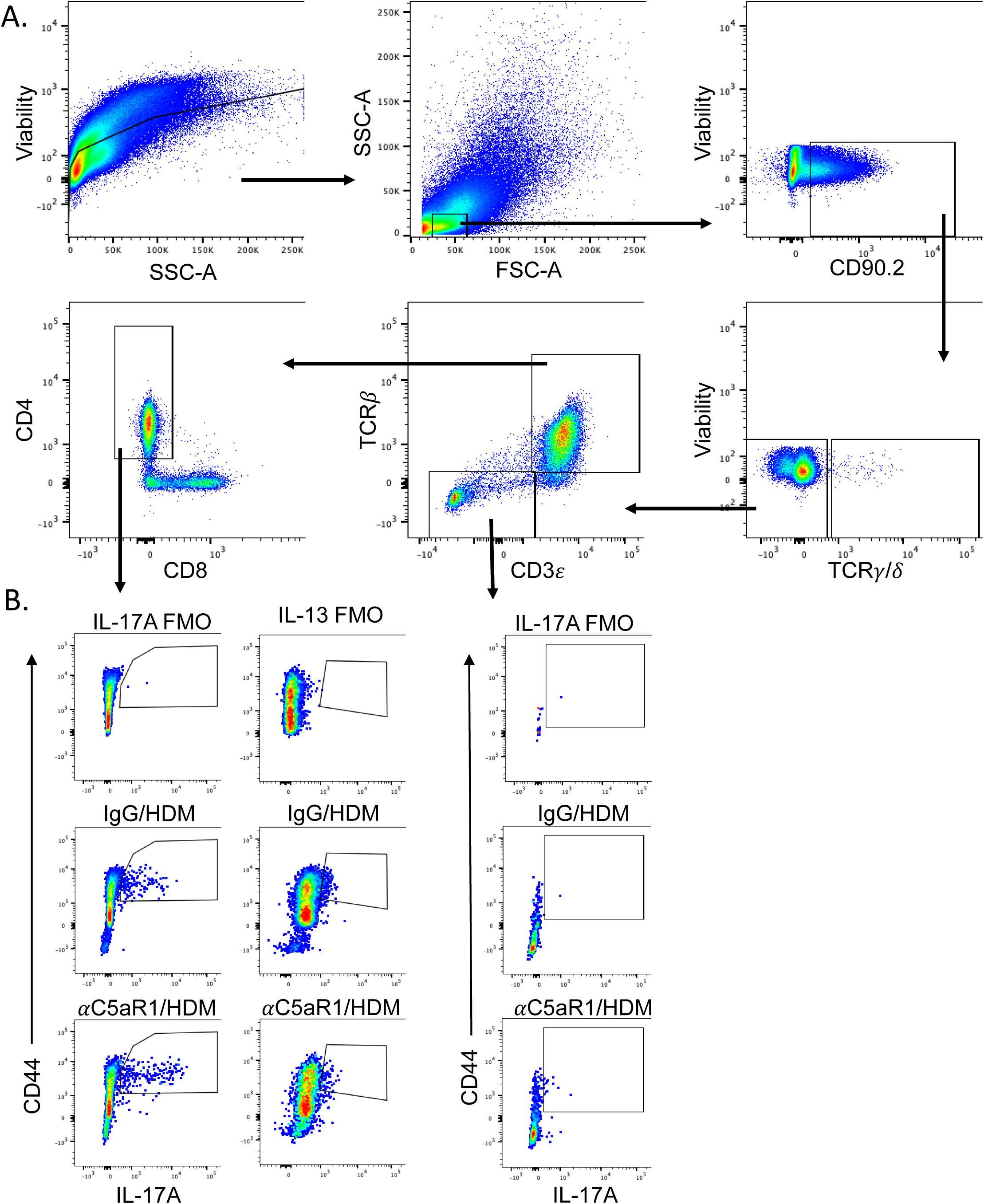
Flow cytometry gating strategy of lung tissue from HDM+ IgG (mild) and HDM+ αC5aR1 (severe) AI mice. (A) Gating strategy to examine IL-17A and IL-13 production by CD4+ T cells or IL-17A production by ILCs. IL-17A (IL-13) expressing CD4+ T cells were identified as CD90.2+,𝛾δTCR-,CD3+TCR𝛽+, CD4+, CD44+IL17A+(IL-13+) live lymphocytes. IL-17A expressing ILCs were identified CD90.2+,𝛾δTCR-,CD3-TCR𝛽-, CD44+IL17A+ live lymphocytes. A representative plot from each treatment group and FMO is shown in panel (B)

**Fig S2:**
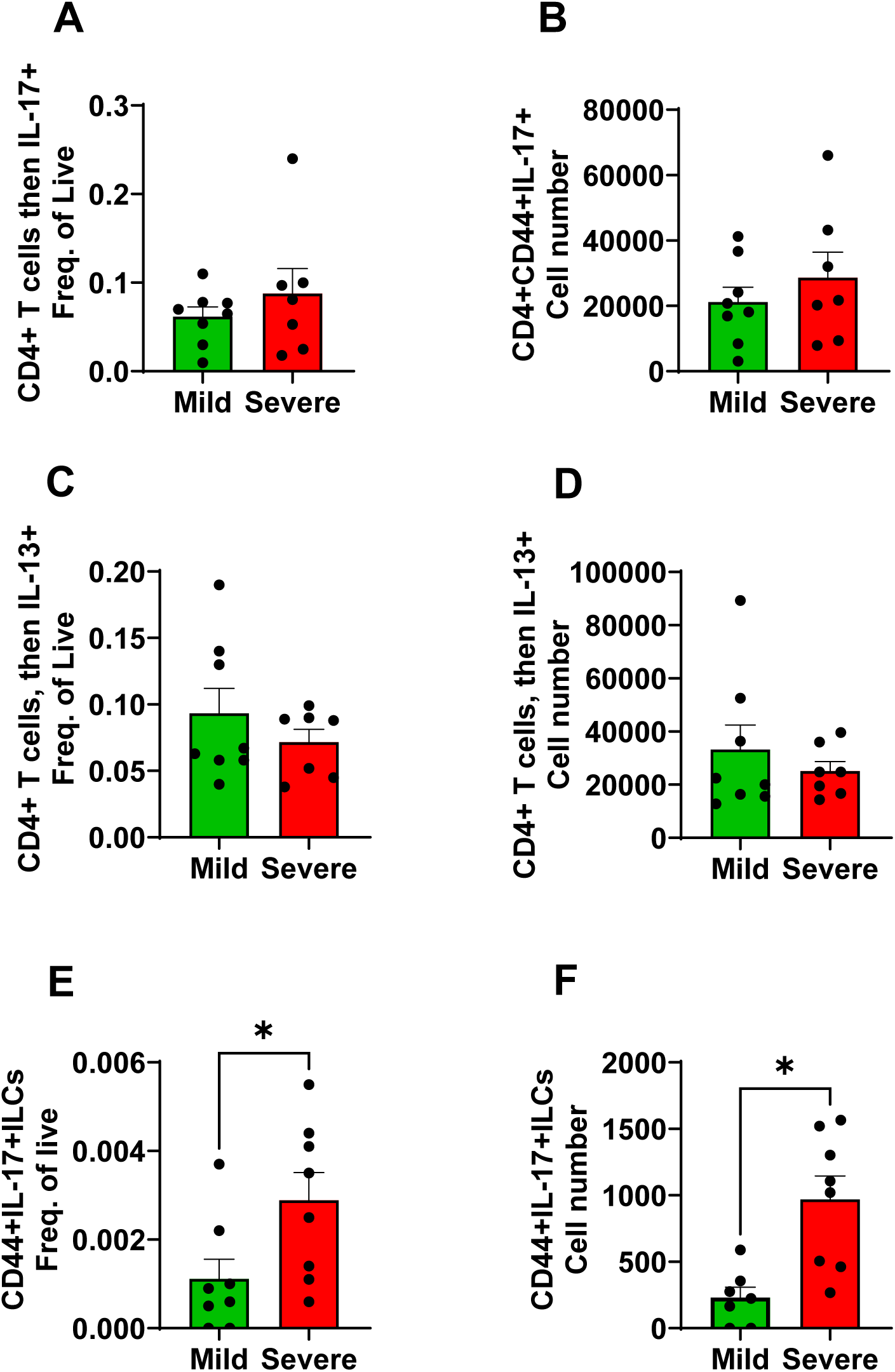
Flow cytometry analysis of lung tissue from HDM+IgG (mild) and HDM+ αC5aR1 (severe) AI mice. The frequency and number of IL-17A (A-B) and IL-13 (C-D) producing CD4+ T cells and IL-17A producing ILCs (E-F) in the lungs of HDM-treated mice that received either IgG antibody (mild) or anti-CD5aR1 antibody (severe). IL-17A (IL-13) expressing CD4+ T cells were identified as CD90.2+,𝛾δTCR-,CD3+TCR𝛽+,CD4+,CD44+IL17A+(IL-13+) live lymphocytes. IL-17A expressing ILCs were identified CD90.2+,𝛾δTCR-,CD3-TCR𝛽-,CD44+IL17A+ live lymphocytes. Data are shown as a frequency of total live cells in each sample as well as the number of cells per lung sample. Significantly higher frequency of total live IL-17A producing lung ILCs (panel E) (unpaired t test t=2.319, df=14, p=0.036) and total number of IL-17A producing lung ILCs (panel F) (unpaired t test t=3.627, df=13, p=0.003) was observed. Mean ±SEM. * p=, students *t*-test, N=8 mice per group.

**Fig S3:**
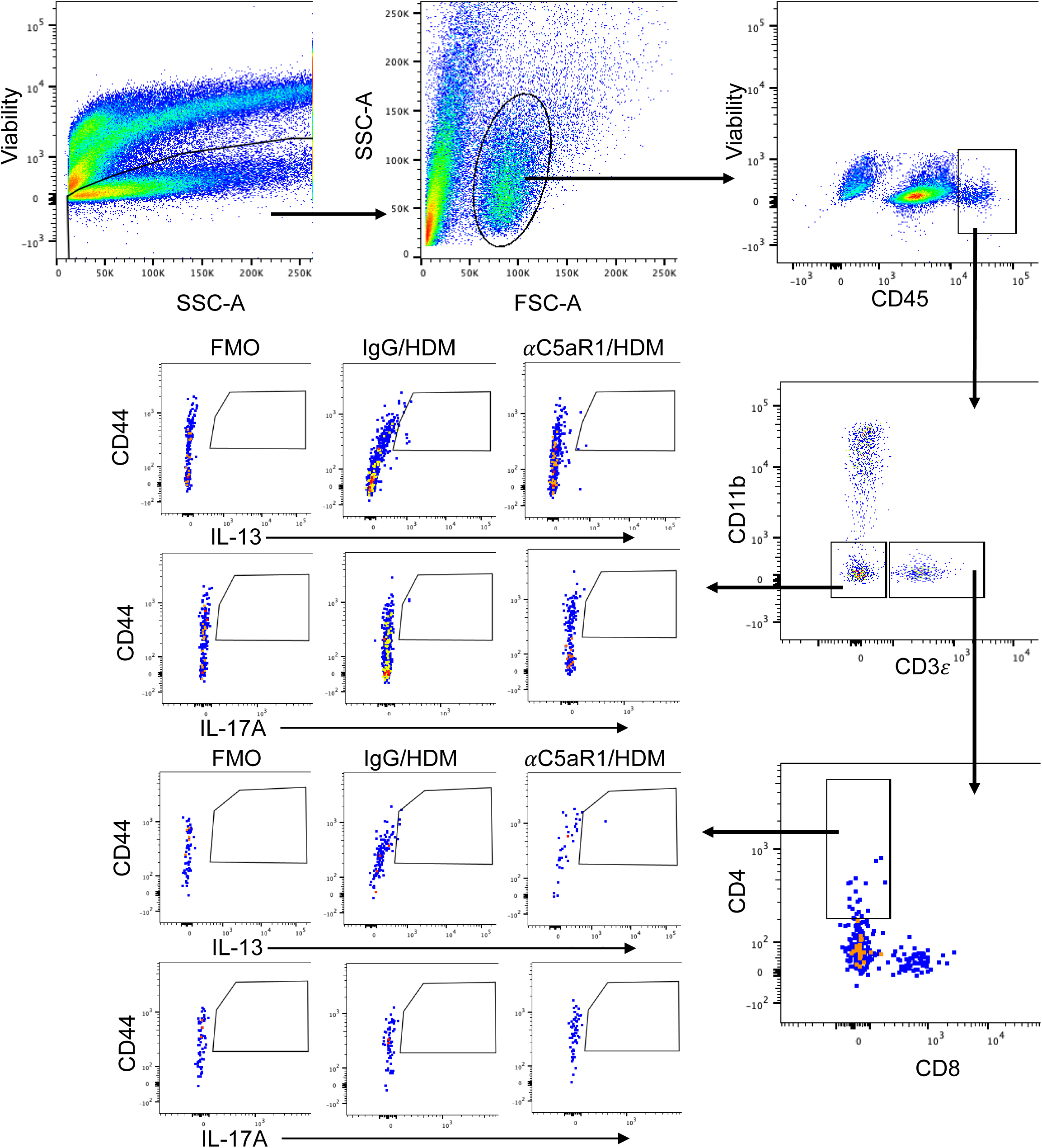
Flow cytometry gating strategy of brain tissue from HDM+ IgG (mild) and HDM+ αC5aR1 (severe) AI mice. Gating strategy to examine cytokine production by infiltrating immune cells in the brain. Infiltrating immune cells were identified as CD45Hi. IL-13- or IL-17A-producing CD4+ T cells were defined as CD3+CD11b-CD44+ while other infiltrating immune cells were identified at CD3-CD11b-CD44+. A representative plot from each treatment group and FMO is shown.

**Fig S4:**
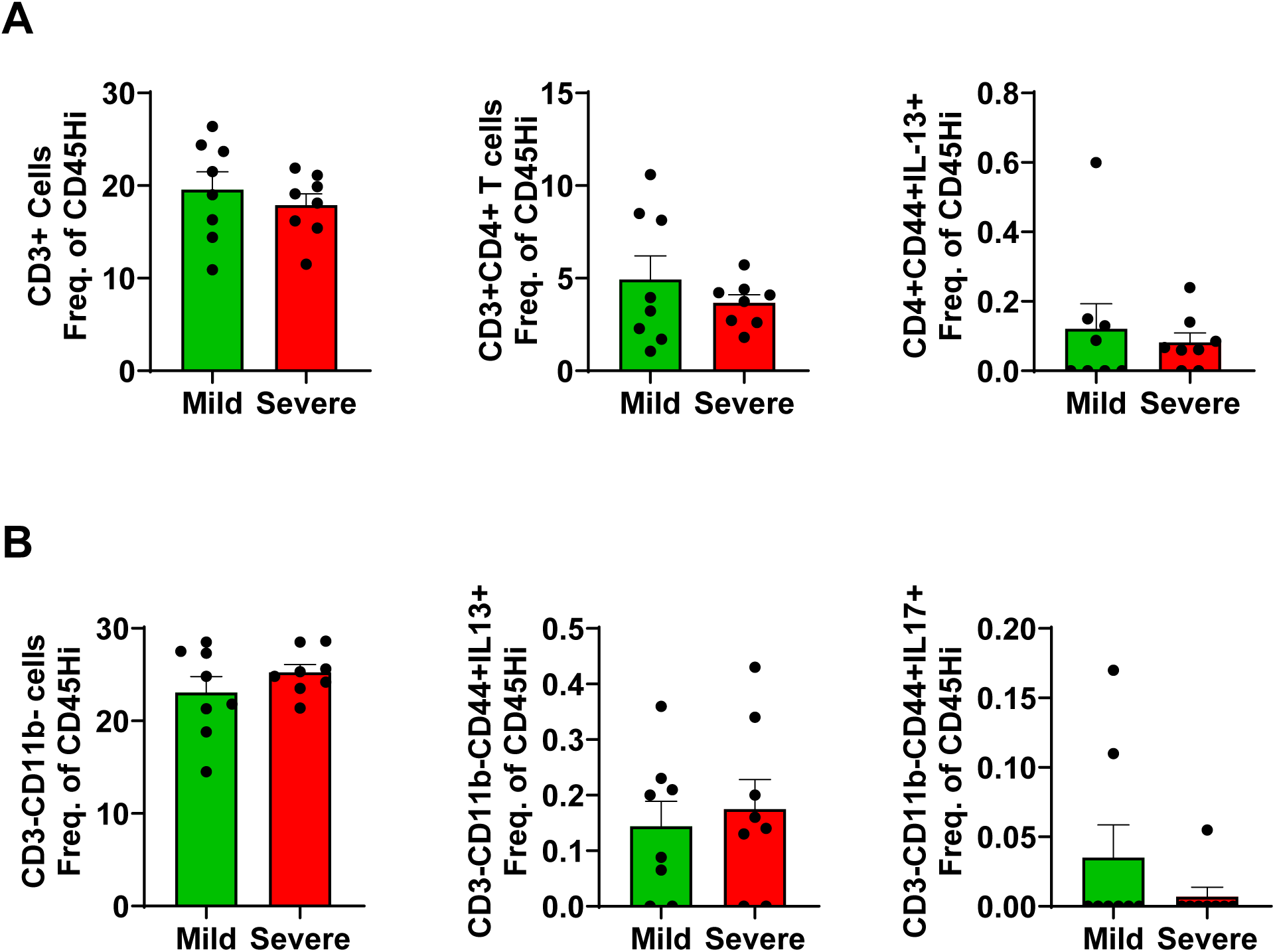
Flow cytometry analysis of brain tissue from HDM+ IgG (mild) and HDM+ αC5aR1 (severe) AI mice. The frequency of infiltrating CD3+CD11b-(A.) and CD3-CD11b-(B.) cells in the brains of HDM-treated mice that received either IgG control antibody or anti-C5aR1 antibody. IL-13-producing CD4+ T cells were defined as CD45HiCD3+CD11b-CD4+CD44+ and IL-13- and IL-17A-producing immune cells were defined at CD45HiCD3-CD11b-CD44+. No differences were observed between treatment groups (p>0.05). Data are shown as a frequency of CD45Hi cells in each sample, ean ± sem. N= 8 mice per group

**Figure S5:**
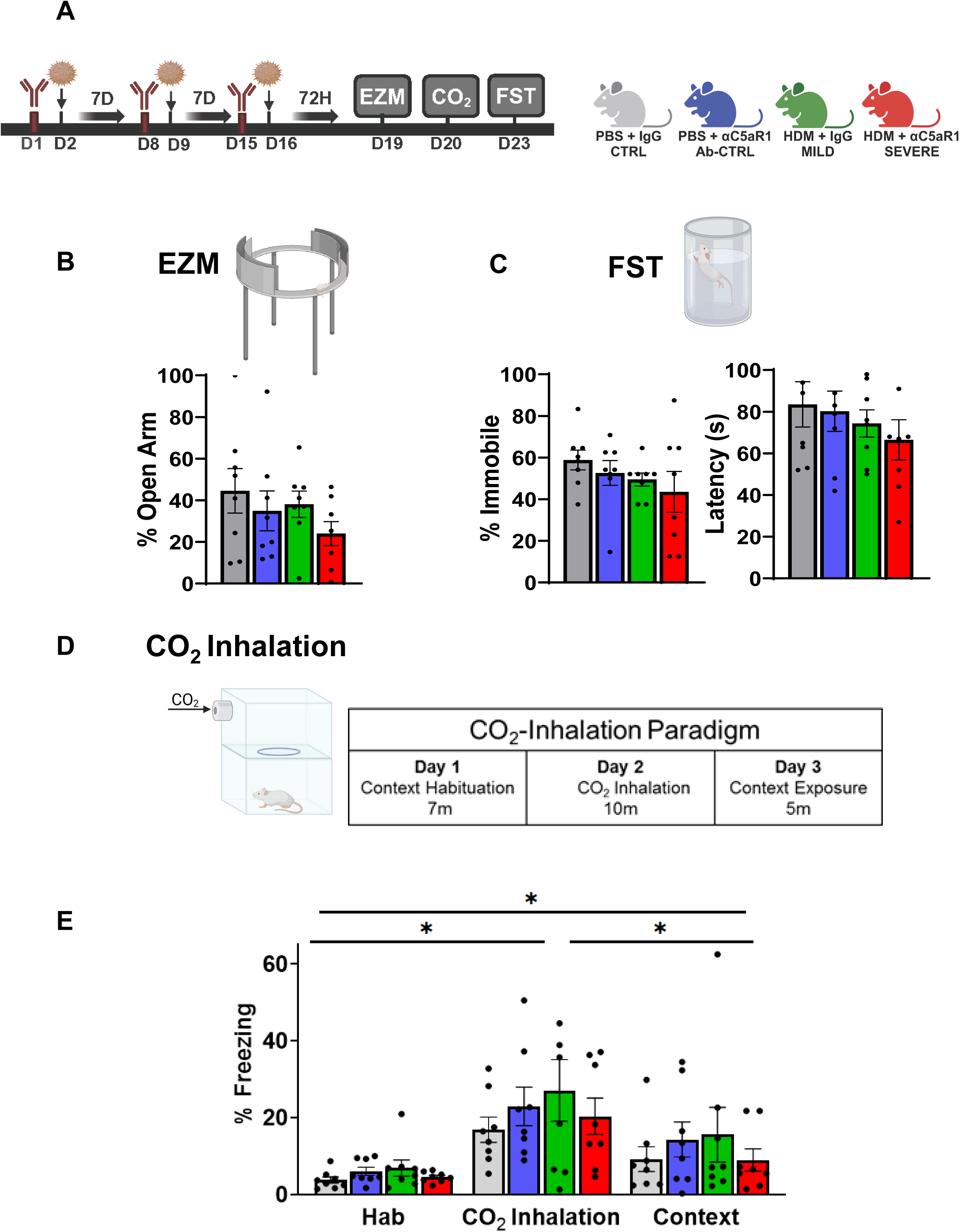
Defensive behaviors to exteroceptive (elevated zero maze, EZM, forced swim test, FST) or interoceptive (CO_2_ inhalation) threat exposures are not altered in severe AI mice. (A) Schematic showing experimental layout and timeline. Mice treatment groups that underwent behavioral testing are shown. (B) Exposure to the EZM showed no significant group differences (open arm time, F_3,28_ = 0.9966; p= 0.4089) (C) No significant group differences were observed in the FST (% immobility, F_3,28_ = 0.9988; p= 0.4079; latency to immobility, F_3,28_ = 0.6089; p = 0.6417) (D) Dual chamber CO_2_-inhalation setup and schematic of the 3-day CO_2_ inhalation-contextual fear paradigm (E) Fear response (% freezing) to interoceptive threat, CO_2_ inhalation and CO_2_-context conditioned fear (CF) revealed increased freezing to CO_2_ inhalation and 24h post CO_2_ context exposure. A 2-way ANOVA revealed a significant effect of exposure day (F _1.719, 48.13_ =30.13, p<0.0001), post hoc analysis revealed significantly higher freezing on CO_2_ and context exposure days relative to habituation [Hab v CO_2_ (p < 0.0001), Hab v CF (p = 0.0044) and CO_2_ vs CF (p < 0.001)]. However, no significant effect of treatment group was observed (F _3,28_ = 0.725, p=0.545). All data are mean ± sem, N=8 mice per group

**Figure S6:**
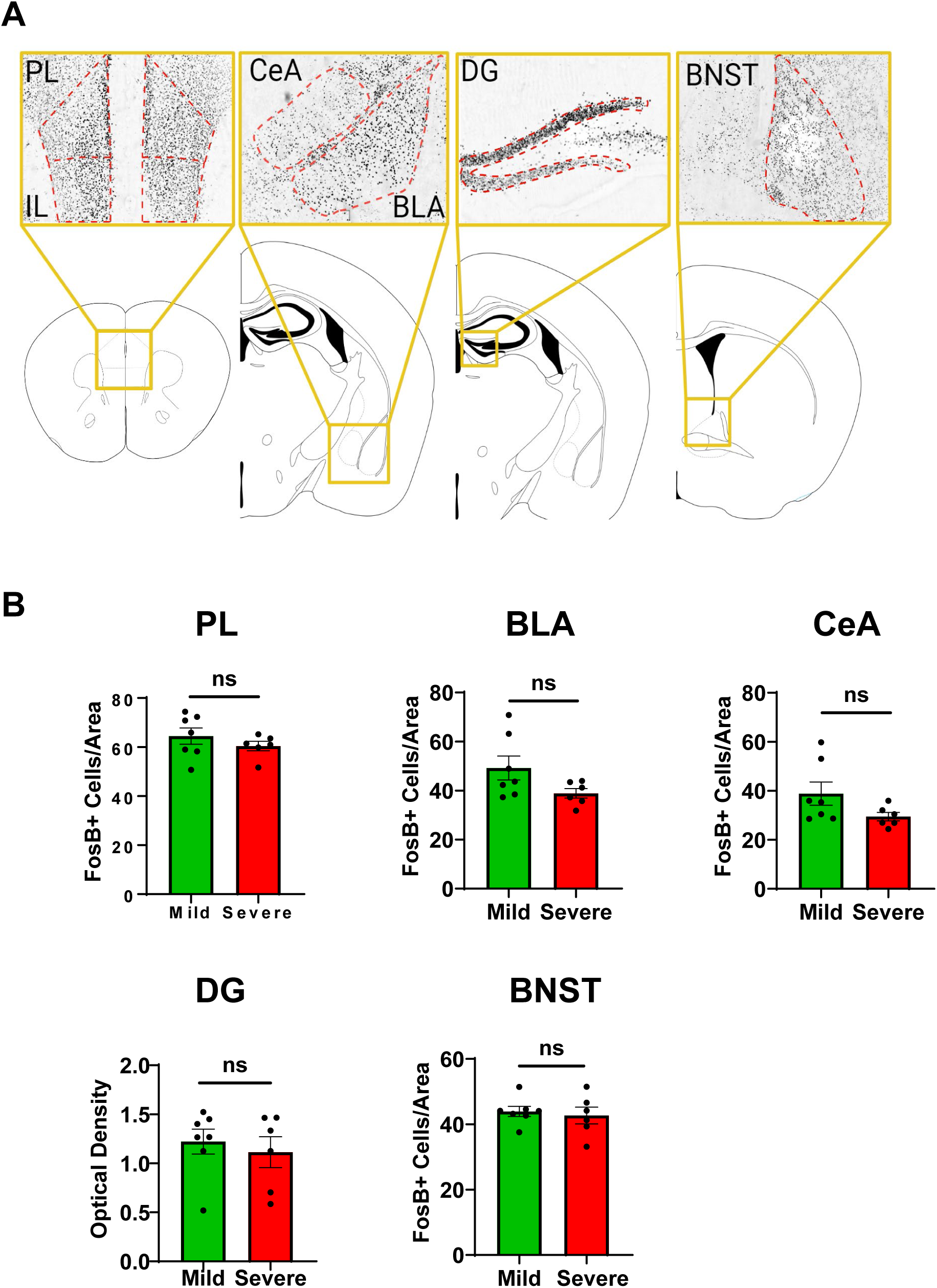
Post behavior ΔFosB cell counts within fear regulatory brain regions. (A) Brain atlas representation of brain areas for ΔFosB quantification: prelimbic cortex (PL), basolateral amygdala (BLA), central nucleus of amygdala (CeA), hippocampal dentate gyrus (DG), and bed nucleus of stria terminalis (BNST) (B) No significant differences were observed in the PL (t_11_ = 0.33; p = 0.330), BLA (t_11_ = 1.85; p = 0.091), CeA (t_11_ = 1.74; p = 0.110), DG (t_11_ = 0.543; p = 0.597) or BNST (t_11_ = 0.435; p = 0.671). All data are Mean ± sem, N= 7 (mild), 6 (severe)

**Figure S7:**
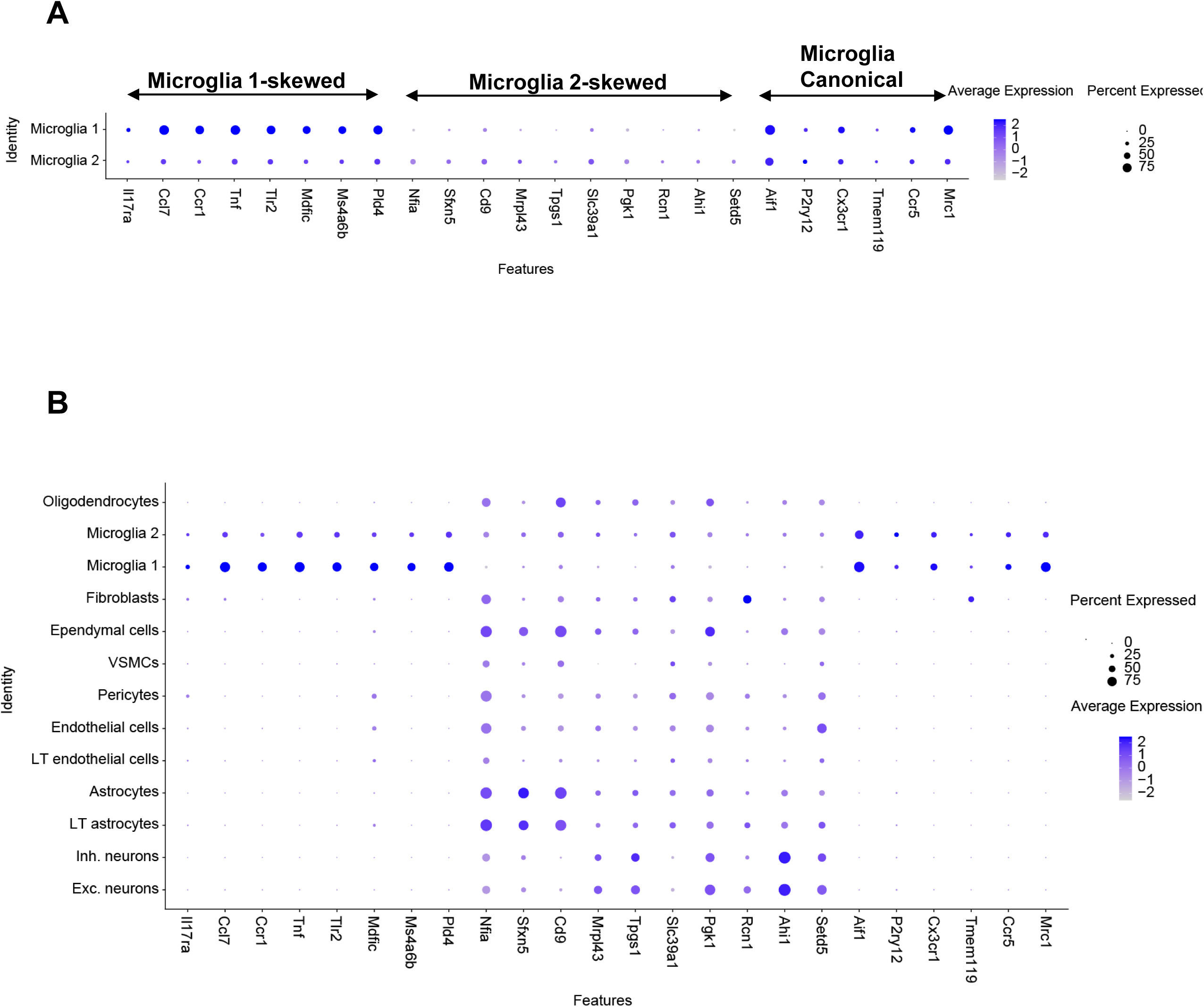
Molecular profile of two transcriptomically distinct microglial populations in the SFO. **A.** Dotplot of cell-type-specific and canonical microglial markers in the two SFO microglial populations. Dot size is proportional to % of cells with transcript count >0, color scale represents z-scored average gene expression (*n* =m7,950 SFO cells). **B.** Dotplot of cell-type-specific and canonical microglial markers in SFO cell classes. Data in **A** and **B** reanalyzed and plotted from a previous study (Pool et al, 2020) [56].

**Figure S8:**
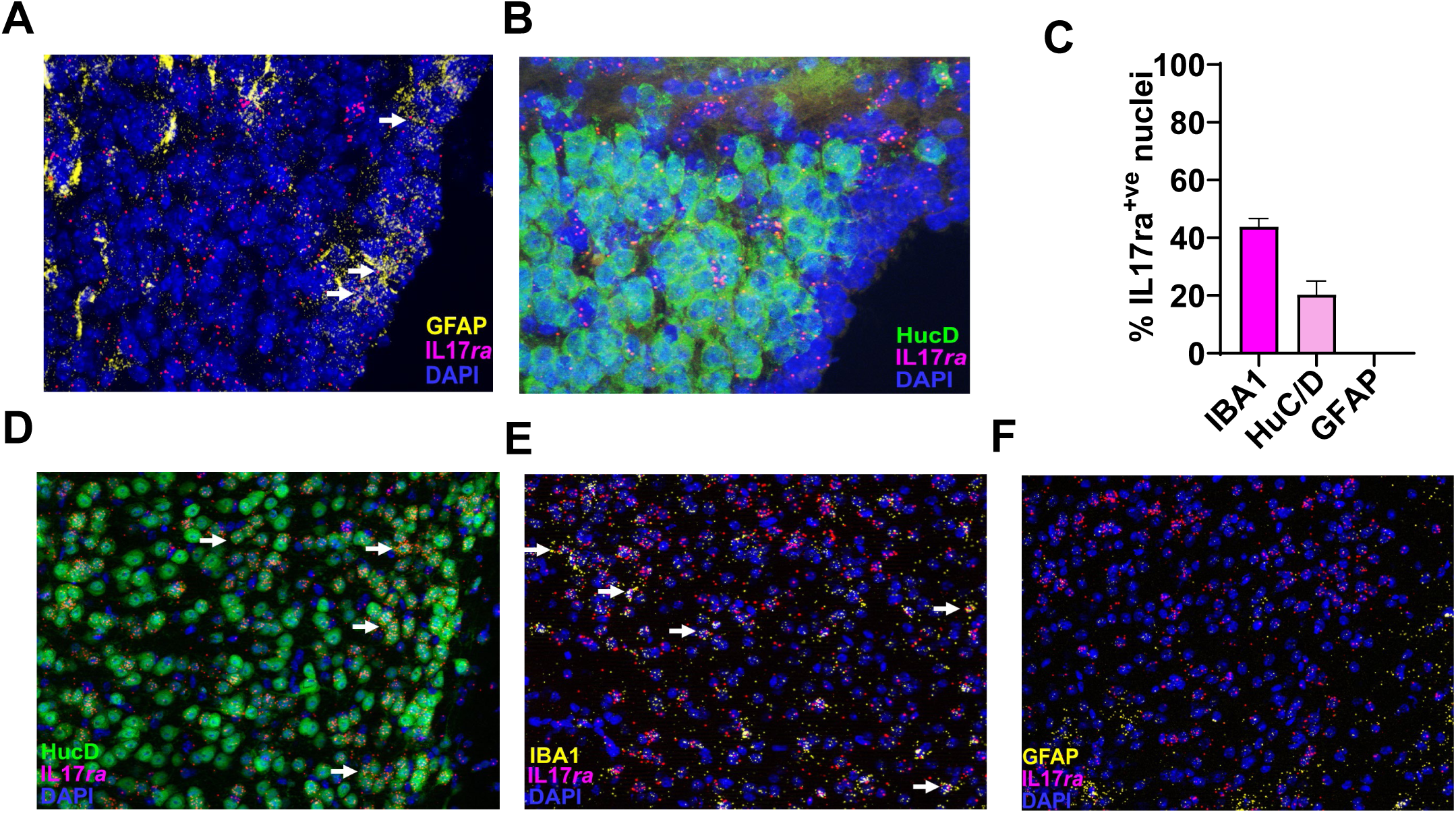
RNAscope in-situ hybridization for cellular expression of *Il17ra* in the SFO and PFC. Representative 20x and 40x images from RNAscope *in-situ* hybridization on SFO slices using probes for *Il17ra* and GFAP (A) and immunohistochemistry with neuronal marker HuC/D (B). Arrows in (A) show scattered co-localization of *Il17ra*^+ve^ puncta (magenta) with GFAP^+ve^ puncta (yellow), DAPI stained cells shown in blue. No co-localization of *Il17ra*^+ve^ puncta (magenta) was evident with HuC/D^+ve^ neurons (green) (B). Graph (C) and Images (D-F) are from *Il17ra* in situ-hybridization performed in PFC slices. Representative 20x images using *Il17ra* probe with neuronal marker HuC/D (D) and IBA1 (E) co-localized with *Il17ra* puncta. Arrows in (D, E) show scattered co-localization of *Il17ra*^+ve^ puncta (magenta) with HuC/D^+ve^ cells (green) and IBA1^+ve^ puncta (yellow) (E). No co-localization was observed with GFAP^+ve^ puncta (yellow) (F). Graph (C) shows percent expression of *Il17ra* with IBA1 (43.8%), HuC/D (20.3%) and GFAP (not detected) within the prefrontal cortex.

